# Tubule extension by plastids is due to increase in bilayer-forming galactolipids at the outer envelope membrane

**DOI:** 10.64898/2026.10.05.756051

**Authors:** Jaideep Mathur, Laura M.J. Currie, Mariel Imada Burnside, Karthika Sriskantharajah, Jose A. Casaretto, Kiah A. Barton, Michael Wozny, Liping Wang, Neeta Mathur

## Abstract

Sporadically, all plastids produce dynamic tubular extensions in response to myriad stresses. Although the use of fluorescent proteins revealed these transient tubules to be stroma-filled and led to their being called stromules, the mechanism underlying tubule extension from a spheroidal plastid remained unknown. As plastids in viridiplantae have two bounding membranes the fluid stroma remains confined within the inner envelope membrane (IEM) while the outer envelope membrane (OEM) around the IEM forms the interface with the rest of the cytoplasm. Here, using a sucrose treatment for inducing tubules in the dark in transgenic Arabidopsis plants expressing different stroma and OEM-targeted fluorescent proteins we carried out comparative molecular-genetic and lipidomics analyses. Our data strongly suggest that plastid tubule formation occurs due to increase in bilayer-forming galactolipids such as digalactosyldiacylglycerol (DGDG) on the OEM. While the dynamic behaviour of stromules occurs within an external encasing OEM, stresses such as phosphate starvation and high light intensity cause the galactolipid-rich OEM-tubule projecting beyond the stromule to be shed into the cytoplasm. OEM shedding from long tubules with extended plastid outreach into the cell suggests their important role in dispersing plastid-derived, membrane-stabilizing, galactolipids to the rest of the cell to augment its resilience to stress.

## Introduction

Plastids in the viridiplantae have an envelope comprising of two membranes, an inner envelope membranes (IEM) that delimits the fluid stroma, and an outer envelope membrane (OEM) that forms the interface between the plastid and the rest of the cytoplasm. All types of plastids have been observed responding to a wide variety of stresses by forming thin, dynamic, several micrometres long, tubular extensions of the envelope that increase plastidial outreach into the cytoplasm (Wildman et al., 1962). The use of a green fluorescent protein highlighted the stroma inside the tubules and led to their being called stromules (Köhler et al. 1997; Köhler and Hanson, 2000). In the years since the initial fluorescent protein-aided discovery of stromules a large number of investigations have established their formation in response to a variety of abiotic and biotic stresses, in diverse cell and tissue types at different stages of development and using experimental systems such as protoplasts, tobacco BY2 cells (Rahpeyma et al., 2026), excised leaf discs and entire plants (Gray et al., 2001; Natesan et al., 2005; Hanson and Conklin, 2020). Notably, while all plastids have the potential to form stromules, at any given time they are extended by only a small sub-population of plastids in a cell, ranging from 0.5 to 35% (Barton et al., 2018). Many of the analyses carried out on extended stromules have been used to furnish information about their alignment with cytoskeletal elements and other membranes (Kwok and Hanson, 2003; Natesan et al., 2009; Schattat et al., 2011a; 2011b; Kumar et al., 2018; Erickson and Schattat, 2018; Meier et al. 2023). Nevertheless, the function of stromules also remains unclear as most studies involving the extensions suggest functions relating to the specific condition under which the stromules were observed. Accordingly, stromules have been implicated in exchange of proteins between plastids (Köhler et al., 1997), in inter-plastid communication (Langeveld et al., 2000), in a plastid density sensing mechanism within the cell (Pyke and Howells, 2002), in defining a chloroplast-nuclear interaction zone (Erickson et al., 2017), in pathways between nuclei and more distant regions of the cell (Kwok and Hanson, 2004), in facilitating retrograde information exchange with the nucleus (Exposito-Rodriguez et al., 2017; Mullineaux et al. 2020), in sensing and dissipating high light induced ROS in chloroplasts (Brunkard et al., 2015), as well as in creating conduits between cells (Krenz et al. 2012). Stromule involvement is also strongly indicated in response to pathogens (Krenz et al., 2012), in innate immunity in plants (Caplan et al. 2015), and in chloroplast degradation and piecemeal autophagy (Ishida et al., 2014).

Despite the large number of investigations focused on stromules an understanding of the exact mechanism whereby a spheroidal plastid produces a long tubule has remained elusive. Considering the fact that a stromule may remain extended for upwards of 30 minutes or display rapid extension and retraction within a couple of minutes, it is unclear how the dynamic behaviour is achieved while maintaining a nearly constant tubule diameter. An issue that may have played a role in limiting our understanding of the dynamic behaviour of stromules appears to lie in assuming that all tubules observed from plastids are stromules. While some reports (e.g. Breuers et al., 2012) have used OEM-targeted fluorescent proteins and refer to the tubules as stromule-like, the majority of observations on stromules have relied on stroma-targeted probes. Observations that would consider that an entire bi-layered tubule would consist of an OEM encasing the stroma-filled, IEM-delimited portion (Fig. 1A), have been singularly missing. While under normal conditions of growth and development the IEM and OEM may remain very close, in the range of 5 to 10 nm from each other (Keegstra et al., 1984; Cline et al., 1985; Cremers et al., 1988; Engel et al., 2015) their proximity and the size of the inter-membrane region (IMR) may not always remain the same. In this context, it is not known whether the spatial relationship between the two membranes remains the same under conditions that overly favour addition of bilayer-forming galactolipids to the OEM.

**Figure 1.**
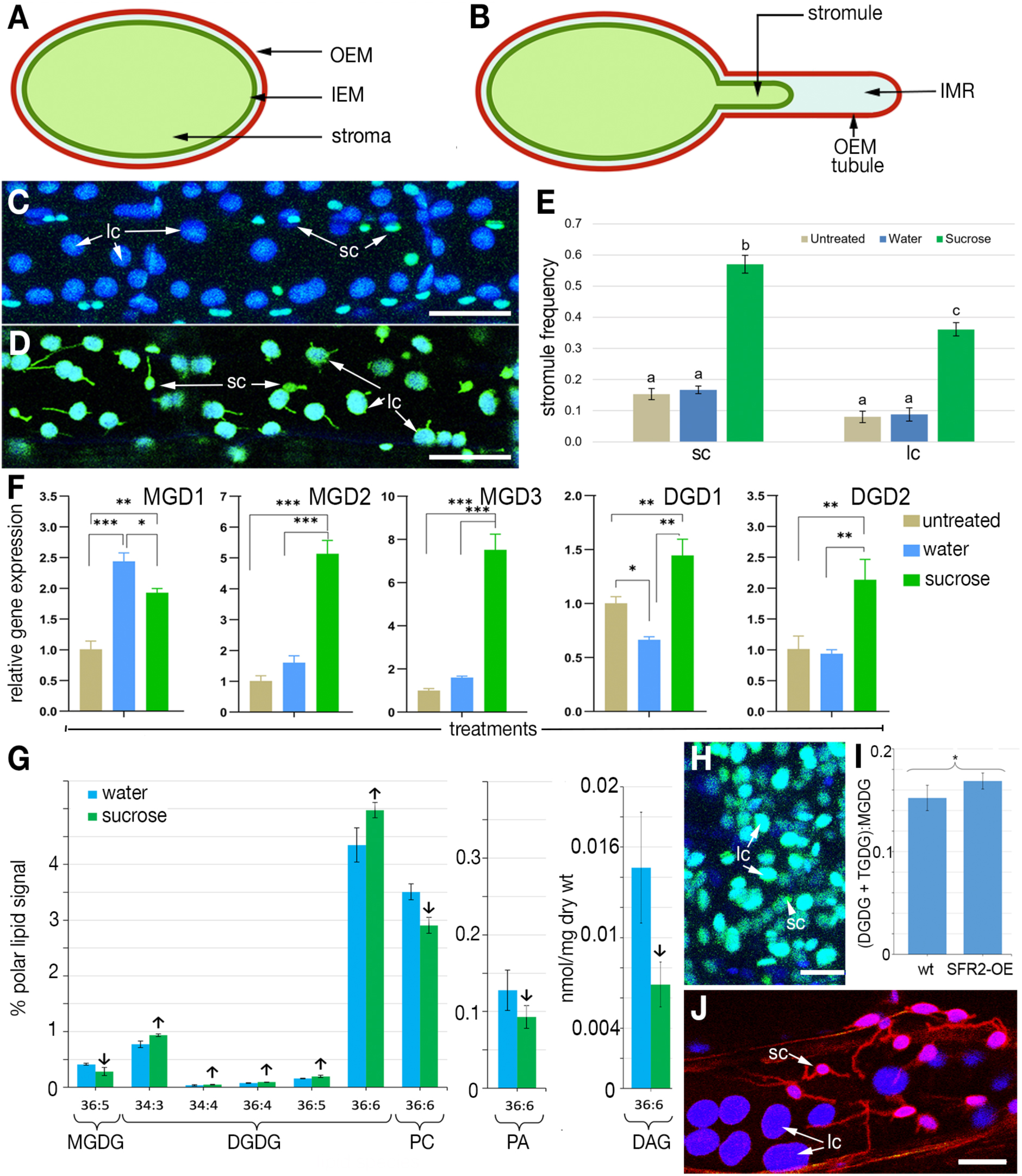
Diagrams, representative images and data supporting links between plastid tubule formation and increase in bilayer-forming galactolipids. A. Diagram showing the spatial relationship between the stroma, confined within the Inner envelope membrane (IEM) and the outer envelope membrane (OEM). B. Diagram forming the hypothetical basis for investigating the altered spatial relationship between the IEM-delimited stroma-filled tubule (stromule), and an elongated OEM-tubule that extends beyond the stromule. The inter-membrane region (IMR) may expand considerably following IEM and OEM separation. C. Chloroplasts in hypocotyl cells of 7-day-old soil grown plants expressing tpFNR-EGFP kept in the dark for 12 hours do not exhibit stromules in small chloroplasts (sc) or large chloroplasts (lc). D. Representative image showing chloroplasts in hypocotyl cells of 7-day-old soil grown plants expressing tpFNR-EGFP that were kept in the dark for 12 hours before treating with 40 mM sucrose for additional 4 hours in the dark exhibiting extensions from both sc and lc (Supplementary Movie 1). E. Stromule frequency is higher in small chloroplasts (sc) than large chloroplasts (lc) after treatment with 40 mM sucrose. Bars indicate the average stromule frequency ± SE of 115 biological replicates with 16 technical replicates each. Varying letters indicate significant differences as determined with a Tukey HSD test at the p < 0.01 level of significance. The water and sucrose treatments formed the basis for subsequent qPCR and lipidomics analyses. F. qPCR-based comparison of expression levels for MGDG and DGDG synthases for plants grown on plates containing ½ MS medium supplemented with 0.75% sucrose and left untreated, water-treated and 40 mM sucrose treated for 4 hours in the dark. G. Lipidomics-based estimation of lipids in plants kept in the dark for 12-hours before being treated in the dark with water and 40 mM sucrose for additional 4-hours. While 36:5 MGDG levels decrease there are significant increases in different DGDG species. Other species of MGDG did not change. Clear decreases were also observed for 36:6 PC, 36:6 PA and 36:6 DAG, implicating the eukaryotic lipid synthesis pathway for the creation of DGDG. Arrows indicate statistically significant differences between treatments as determined by Student’s t-tests (P < 0.05). H. Representative image showing that stromules are not formed in a *dgd2dgd1* double mutant after a 4-hour treatment with 40 mM sucrose. Chloroplasts in plants expressing tpFNR-EGFP in wild type background (B) extend stromules at high frequency under the sucrose treatment. I. The SFR2-mRFP transgenic line #14 shows the OEM of small epidermal chloroplasts (sc) in hypocotyl cells of 7 -10 days old plants extending tubules exclusively after 12 hours of darkness. By comparison, the OEM of large, sub-epidermal chloroplasts (lc) is also fluorescently highlighted but no tubular extensions are observed. J. Average ratio between the lipid content of bilayer-stabilizing galactolipids, (DGDG and TGDG), and bilayer-disrupting lipid MGDG in the *A. thaliana* SFR2-mRFP OE #14 with respect to the wildtype. Average ± standard deviation calculated from five independent samples. Asterisks indicate statistically significant differences between lines as determined by a Student’s t-test (P < 0.05). Size bars in C, D =25; H, J =10 µm. Abbreviations: DAG – diacylglycerol; DGD1; -DGD2 - di-galactosyldiacylglycerol synthase 1 and 2, respectively; DGDG – di-galactosyldiacylglycerol; IEM - inner envelope membrane ; IMR – inter membrane region; lc – large sub-epidermal chloroplasts; MGD1- MGD2- MGD3 - mono-galatosyldiacylglycerol synthase1;-2;-3, respectively; MGDG – mono-galatosyldiacylglycerol; OEM -outer envelope membrane; PA – phosphatidic acid; PC – phosphotidylcholine;; sc – small epidermal chloroplasts; SFR2 – SENSITIVE TO FREEZING 2; TGDG – tri-galactosyldiacylglycerol.

Understanding stromule dynamics within the confines of an encasing OEM-tubule also becomes specially relevant when one considers that the lipid composition of the IEM and OEM differ considerably. Plastid membranes in green plants primarily contain galactolipids (Benning and Ohta, 2005) that make them different and relatively more stable compared to phospholipid membranes in a cell (Róg et al., 2007). Of the two major plastid galactolipids, mono-galactolsyl-diacylglycerol (MGDG) and digalactolsyl-diacylglycerol (DGDG), the former is relatively more abundant on the IEM, compared to the bilayer-forming DGDG (Froehlich et al., 2001; Kelly and Dörmann, 2002; Jouhet 2013; Hölzl and Dörmann 2019). By contrast, the OEM has a high DGDG to MGDG ratio (Douce and Joyard, 1990; Block et al., 2007; Benning 2009) which results in its greater bilayer-forming capability (Cline and Keegstra, 1983). In addition to the UDP-galactose dependent MGD and DGD synthases an additional, UDP-galactose independent, processive galactolipid:galactolipid galactosyltransferase (GGGT) protein-based mechanism (Dorne et al., 1982; Cline and Keegstra, 1983; Heemskerk et al., 1986) also helps in OEM stabilization (Moellering and Benning, 2011; Roston et al., 2014). Encoded by SFR2 (SENSITIVE TO FREEZING 2; At3g06510; Moellering et al., 2010) the GGGT activity is triggered naturally under relatively harsh stresses such as exposure to ozone (Sakaki et al., 1990a;1990b), desiccation (Gasulla et al., 2013) drought, salinity (Wang et al. 2016), freezing (Moellering et al., 2010), and wounding (Vu et al., 2014; 2015) and leads to the formation of bilayer-stabilizing oligogalactolipids (OGDG) such as tri- and tetra-galactolsyldiacylglycerol (Mugume et al., 2026).

Based on the well-established compositional differences between the two membranes of the plastid envelope (Douce and Joyard, 1990; Benning and Ohta, 2005), with the OEM being richer in bilayer-forming lipids such as DGDG and OGDG we hypothesized that a further increase in bilayer-forming galactolipids would specifically favour OEM stabilization and its extension beyond the IEM. We speculated that a long, stabilized OEM-tubule could accommodate both stromule form and its dynamic behaviour.

A major limitation in investigating plastid tubules has been their sporadic occurrence at a low frequency. Therefore, we used the established tpFNR-EGFP stroma marker line ((Marques et al., 2003; Schattat and Klösgen, 2011) to identify two contrasting conditions; one, twelve hours of darkness, where the number of stromules reached a minimum (Barton et al., 2018), and the second, treating plants with 40 mM sucrose where stromule frequency has been shown to increase significantly (Schattat and Klösgen, 2011). Analyzing plants subjected to these two conditions with a range of molecular-genetic, lipidomics and cell-biological techniques set the stage for further investigations using a set of double transgenic Arabidopsis lines where the stroma and the OEM are highlighted by spectrally well-separated fluorescent proteins. The simultaneous visualization of a stromule and its encasing OEM-tubule has led to significant insights on the mechanism of plastid tubule formation and strongly suggests tubule involvement in dispersing plastid derived, membrane-stabilizing galactolipids within the plant cell.

## Materials and methods

### Gene cloning

A modified pCAMBIA 1300 vector (Roberts et al., 2000) with CaMV35S promoter was used to create three separate cassettes having either EGFP (Haseloff et al., 1997), mRFP (Campbell et al. 2002) or mEosFP (Wiedenmann et al., 2004) for expression of fluorescent fusion proteins. Gene specific primers used are listed in Supplementary Table 2.

### Transient overexpression

Transient expression over 36 hours was used to check all fusion constructs through *Agrobacterium tumefaciens* strain strain GV3101::pMP90 (Koncz and Schell, 1986) infiltration of leaves of *Nicotiana benthamiana*.

### Creation of transgenic plants

The creation of transgenic lines tpFNR-EGFP and tpFNR-mEosFP (Schattat et al., 2012a), SFR2-mRFP (Mathur et al., 2022), RFP-HDEL (Sinclair et al., 2009) has been reported earlier. All stable lines used (Table 2) were created in the *Arabidopsis thaliana* ecotype Columbia and in the *hps1* (Lei and Liu, 2011), *dgd2dgd1*, *pah1pah2* (Eastmond et al., 2010) and *tgd1*(Xu et al., 2003) mutants by floral dipping (Clough and Bent 1998). Double and triple transgenic lines were created either by direct *Agrobacterium* mediated transformation or through genetic crossing. The majority of observations reported here were taken on 8 to 10 days old intact seedlings following their germination in soil boxes under 110 ±10 μmol m^-2^ s^-1^ photosynthetically active radiation (PAR) and a 16/8-hour light/dark cycle at 21 ± 2 °C. Some independent experiments involved seed gemination and subsequent growth on ½ MS (Murashige and Skoog, 1962) medium containing 0.75% or 1% sucrose and are mentioned as such in text.

### Gene expression analysis

Total RNA was isolated from 100 mg of whole seedlings grown as indicated above using the Total Plant/Fungal RNA Isolation Kit (Norgen Biotek Corp). RNA was treated by DNase I (Promega) and then 1 μg of DNA-free total RNA was reverse transcribed into cDNA using the LunaScript® RT SuperMix Kit (New England Biolabs) according to the manufacturer’s protocol. Real-time qPCR reactions (20 µl) were performed using the Luna® Universal qPCR Master Mix (New England Biolabs) on the ABI7300 system (Applied Biosystems) following the manufacturer’s protocol. The PCR program adjusted for the amplification of the target fragments consisted of one cycle at 95°C for 2 min, and 40 cycles of 95°C for 15 s and 60°C for 40 s. The gene-specific primers were designed with the software Primer3 and are listed in Supplementary Table 2. Relative expression was calculated as fold change using 3 biological and 2 technical replicates of each treatment based on untreated control using the 2^−ΔΔCt^ method (Livak and Schmittgen 2001). The At*Actin7* and At*Tubulin7* genes were used as the internal reference for normalization.

### Treatment of plants to assess stromule responses

Plants were taken at the end of night for treatment, unless a period of light is specified for an experiment. Treatment with solutions was carried out by first briefly immersing the whole plant or seedling, unless otherwise specified, in the solution. For assessment in the hypocotyl of 7-day-old seedlings, the seedlings were then floated such that the hypocotyl was submerged but the cotyledons were floating on the surface. The plants were then left in the solutions in the dark at room temperature for 4 h, unless specified otherwise for an experiment. All solutions of sugars and sodium chloride were prepared in deionized water and made fresh for each experiment (Sucrose – BioBasic SB0498, D-Glucose – Sigma G8270; NaCl – SOD001.1 Bioshop).

### Stress treatments

All treatments were carried out on tpFNR-EGFP plants. High temperature stress: Plants grown for 8 days in soil-boxes at 21°C were transferred for 4 hours to an incubator maintained for 31°C, before observing. Osmotic stress: plants dipped for 4 hours in 200 mM Mannitol (M562-Phytotechnology Labs) dissolved in water. Dehydration: Plants taken from soil and left for 2 hours on an open lab bench before comparing to plants in soil boxes. Wounding: Following the leaf crimping method of Vu et al., (2014). Wounded leaves observed with 30 minutes. Ozone treatment: An Ozone generator (CE RoHS model GL-3189A) was used to bubble the gas through de-ionized water for 10 minutes before treating 8 days old seedlings with the ozonated water for time ranging between 2-8 minutes.

### Nutrient deprivation experiments

These were carried out using individual solutions of the macronutrients (final concentrations: 1650 g/L NH4NO3; 332.2 g/L CaCl2; 180.7 g/L MgSO4; 1900 g/L KNO3) supplemented with a micronutrient solution (Sigma M0529) and a solution of Gamborg’s vitamins (Sigma G1019) according to manufacturer’s instructions. For the phosphate-containing control medium, 170 g/L of KH2PO4 was added. Nitrogen deprivation media were prepared using MS media without nitrogen (Caisson MSP21), and 2400 g/L NH4NO3 was added to control media. For all three nutrient deprivation experiments, control and nutrient lacking media were prepared with 0% and with 3% sucrose and were solidified with 1% tissue culture grade agar (Caisson A038). Seeds were plated on MS media with 3% sucrose and were grown for 5 days before transfer to the media prepared for the nutrient deprivation experiments (Nutrient 3% sucrose; Nutrient 0% sucrose; No nutrient 3% sucrose; No nutrient 0% sucrose). 7 days after transfer, seedlings were imaged for stromule frequencies and tissue was harvested for carbohydrate estimates.

### Measurement of total soluble carbohydrates

The soluble carbohydrate measurement procedure was adapted from two existing protocols (DuBois et al., 1956; Buysse and Merckx, 1993). Details provided in Supplementary information.

### Acyl-lipid / lipidomics methods

Lipids were extracted following standard protocols adapted from Shiva et al., (2018). For fatty acid analysis total lipids were stored in a final volume of 50 µL of Chloroform, flushed with nitrogen gas, capped tightly, and stored at -80 °C until the FAME reaction was carried out according to Wang and Benning, (2011). GC analysis was carried out utilizing an Agilent 8890 GC System equipped with the Agilent 7693A autosampler. The sum of the mMol/g FW of all lipid classes of each lipid species was calculated for each replicate, then average and standard deviation were plotted on a bar graph. Lipidomics analyses were carried out using direct-infusion electrospray ionization tandem mass spectrometry (ESI-MS) at the Kansas Lipidomics Research Center Analytical Laboratory < https://www.k-state.edu/lipid/ > according to Shiva et al., (2013).

### Thin layer chromatography (TLC) based analysis

Plants were grown either on soil or ½ MS ¼ sucrose plates with or without sucrose and phosphate as noted, under white light with an 18-hr light, 6-hr dark photoperiod for 10 days before extraction of total lipids according to Shiva et al.,(2018) utilizing a Chloroform: Methanol: Isopropanol: Water ratio of (30:51.5: 25: 3.5) (v/v/v/v) with 0.01% BHT and (KCl:H2O) (1:5) (v/v) for phase separation of lipids. The final ratio is (3.2: 2) for (Chloroform: Methanol: Isopropanol: Water) : (KCl:H2O). Lipids were separated on a TLC plate as described by Wang and Benning (2011) utilizing a glass, silica-coated plate pre treated with 0.15M Ammonium sulfate and baked for 3 hours at 120° C. A developing solvent of 91 mL Acetone : 30 mL toluene : 7.5 mL water was used for a running time of 1 hr. The plate was stained according to Churchward et al., (2008) by spraying with 10% Cupric acid and 10% phosphoric acid and charred at 180 ° C for 10 minutes.

### Electrolyte Leakage assay

The percent electrolyte leakage technique was adapted from Bajji et al.,(2001) and Hezema et al., (2021). Briefly,130 mg of whole seedlings were subjected to pre-treatments, rinsed with miliQ water, then transferred to 50 mL Falcon tubes with 25 mL of miliQ water and agitated at 100 rpm for 24 hours. The initial electrolyte conductivity (IEC) was measured with an electrolyte conductivity meter (Corning, Checkmate90 field system, 475627). To measure total electrolyte conductivity (TEC), tubes were frozen solid at -20 °C freezer for 24 hrs then thawed to 21 °C, before measuring the TEC. To calculate % electrolyte leakage, the formula (IEC/ TEC)* 100. Twelve replicates were done per treatment for statistical analysis.

### Transmission electron microscopy

Plants were grown for 10 days under normal light conditions on ½ MS medium with 0.75% sucrose and the tissue harvested immediately before submerging in Karnovsky’s fixative in phosphate buffer, stained in 1% osmium tetroxide solution (in 0.2 M, pH 7.5 phosphate buffer) for 2 hours, dehydrated through a graded ethanol series, and embedded in Spurs resin. The 88 nm sections, mounted on nickel grids and stained with 3% lead citrate and 2% uranyl acetate in 75% ethanol, were imaged using a 200 kV FEI Tecnai G2 F20 TEM set to a high tension of 120 kV. Images were acquired at 4096 x 6096 pixels, at a magnification between 8000x,12000x, 20000x and a maximum of 50000x.

### Confocal microscopy and time-lapse imaging

All fluorescence imaging was carried out on entire plants mounted in milliQ water on depression slides, using a 40X HCX APO water immersion lens (N.A -0.80). A Leica TCS-SP5 confocal microscopy system equipped with a 488 nm Ar laser and a 543 nm HeNe Laser. GFP fluorescence was acquired between 500-520 nm, RFP fluorescence between 570-620 nm, YFP fluorescence between 530-550 nm and chlorophyll autofluorescence between 660-750 nm. All confocal images were acquired at a resolution of 1024 x 512 or 1024 x 1024 pixels. Three-dimensional (xyz) stacks were collected with a step-size of 0.99 μm and successive frames in x,y,t time lapse series were 1.935 s apart. Images were processed using proprietary Leica (LSM SPII) software, ImageJ and Adobe Photoshop CS6. Quick time 7 was used for assembling and annotating time-lapse movies.

Probes expressing mEosFP (Wiedenmann et al., 2004) were photo-converted from green to red manually using a mercury lamp and the Leica fluorescence filter set D (Excitation 355 to 425 nm) by controlling the diaphragm as described earlier (Mathur et al., 2010; Schattat et al., 2012a). The epifluorescence setup consisted of a Leica DM-6000CS microscope. The light source for photoconversion was an HBO 100 W/2 Mercury Short Arc lamp and the Leica fluorescence filter set D (excitation filter, 355 to 425 nm; dichromatic mirror, 455 nm; suppression filter, long-pass 470 nm).

### Image and data processing

ImageJ software (http://imagej.nih.gov/ij) was used to assimilate data from the CLSM images acquired as Leica (lei.) files. The circularity tool (using the formula: circularity = 4pi(area/perimeter^2^) was used to measure plastid roundness. The freehand line and freehand selection plugins were used for measure stromule lengths, stromule distances from the punctae and the area of punctae and plastids. Extension and stromule frequencies were transformed using the arcsin function for statistical analysis to normalize the data, as outlined by Schattat and Klösgen (2011). The function specifically used ‘ASIN(√(chloroplasts with stromules/total chloroplasts’).

Post acquisition image analysis and histograms were prepared using line segmentation and shape tools for applying the Interactive 3D surface plot (v2.4.1) plugin in imageJ (v2.0.0-rc-65/1.52p). The corrected total cell fluorescence (CTCF) for estimating vesicles in the cytoplasm was calculated using ImageJ software and following Regnier et al., (2024).

### Statistical methods

Two-tailed Student’s t-tests were used to see if there was a statistical difference between two treatments within a plant line or to compare different lines to the same treatment. When comparing the variance between every possible combination of treatment and plant lines, a Tukey’s Honestly Significant Difference (HSD) test was used. Letters above histograms indicate significant differences as determined by Tukey’s HSD method (p<0.001).

Microsoft Excel was used to calculate values such as averages, standard deviations, standard errors, F-tests, and two-tailed t-tests while R statistical analysis software was used to calculate Tukey’s HSD tests.

## RESULTS

### Sucrose availability increases tubular plastid extensions and bilayer-forming DGDG

Based on literature, a diagrammatic depiction (Fig. 1A, B) presents the underlying basis for our subsequent investigations. The plastid envelope comprises of two membranes; an inner envelope membrane (IEM) that delimits the stroma, and an outer envelope membrane (OEM) that surrounds the IEM (Fig. 1A). As shown in Fig. 1B, a stroma-filled tubule (stromule) lies inside an encasing OEM-tubule and is separated from it by an inter-membrane region (IMR). In subsequent usage we have used the word tubule to depict a tube like form, and distinguish it from the word ‘ stromule ’, that by definition refers only to a stroma-filled tubule (Köhler and Hanson, 2000) and lies inside the OEM. We hypothesized that under certain conditions the OEM extends independently from the IEM.

Chloroplasts in Arabidopsis hypocotyl cells are maintained in two sizes, with epidermal cells having characteristically small chloroplasts (sc), about half the size of chloroplasts in sub-epidermal layers (lc) (Barton et al., 2016; 2018; Fig. 1 C). Under normal light conditions, stromules were readily observed from sc but observed only occasionally from the lc. Using an Arabidopsis line expressing a stroma-targeted tpFNR-EGFP (Marques et al.,2003; Schattat et al. 2011a) we observed that after being kept in darkness for 12 hours, less than 2% of sc in hypocotyl cells of 8-to-10-day old plants exhibited stromules (Fig. 1C). However, following an earlier established procedure (Schattat and Klösgen, 2011; Barton et al., 2018) treating the dark-grown plants with 40 mM sucrose for an additional 4 hours in the dark increased the incidence of stromule formation considerably, with up to 29.1 ± 5.5% of sc and 13 ± 3.0% of sub-epidermal lc exhibiting them (Fig. 1D, E; Movie S1). The increased stromule numbers positively correlated with higher total soluble carbohydrates in the 40 mM sucrose treated plants (Fig. S1). Notably, the sucrose treatment resulted in stromule formation from both sc and lc (Fig.1E), which was different from the usual response of lc to light. Since the treatments were carried out in complete darkness it was concluded that stromule formation triggered in all chloroplasts under sucrose availability constitutes a heterotrophic response.

Further support for the sucrose playing an important role in stromule formation came from observations that in comparison to soil-grown plants, with an average of 1.5 ± 0.4% of sc exhibiting stromules at the end of the night, all plastids in tpFNR-EGFP plants grown on half-strength Murashige and Skoog (1962) basal medium supplemented with 1 % sucrose exhibited a higher number of stromules, ranging from 25.3 ± 12.0% of sc and 7.4 ± 2.5% of lc. Since Arabidopsis plants grown on sucrose-supplemented media have been shown to have increased expression of eukaryotic galactolipid synthases including type-B MGDG, and DGDG synthases (Lei et al., 2011; Murakawa et al., 2014) a possible correlation between galactolipids and stromule formation was suggested. Indeed, our qPCR of water and sucrose treated 8-10-day old tpFNR-EGFP seedlings with minimal, and high stromule numbers, respectively, showed significantly increased transcript levels for MGD2, MGD3, DGD1 (Fig.1F). Notably the galactolipid synthases upregulated under 40 mM sucrose treatment contribute to the eukaryotic lipid biosynthesis pathway on the OEM where they have access to eukaryotic diacylglycerol (eDAG) and phosphatidylcholine (PC) substrates (Browse et al., 1986; Ohlrogge and Browse, 1995).

Although our 40 mM sucrose treatment in the dark did not increase fatty acid synthesis in rosette leaves (Fig. S1A), comparative lipidomics of the water and sucrose treated plants showed a decrease in 36:5 MGDG and an increase in amounts of several DGDG species, including 36:5 and 36:6 DGDG (Fig.1G). Concomitantly there was a significant decrease in phosphatidylcholine (PC) 36:6 (Fig. 1G). Notably the galactosylation of phosphatidylcholine (PC)-derived eDAG by UDP-galactose is an integral part of the eukaryotic galactolipid biosynthesis machinery (Ohlrogge and Browse, 1995; Karki et al., 2019) that also involves non-bilayer forming lipids such as PA, DAG and MGDG for the final formation of lamellar (L_a_) phase, bilayer-forming galactolipid DGDG (Jouhet et al., 2003; Botella et al., 2017; Karki et al. 2019). Thus the lower amounts of 36:6 PC, 36:6 PA, 36:5, 36:6 DAG and 36:5 MGDG (Fig. 1G and Fig. S2) with concomitant increase in 36:5 and 36:6 DGDG in our lipidomics analysis further affirmed involvement of the eukaryotic galactolipid synthesis pathway for stabilizing the OEM under sucrose abundance.

Our RNA expression and lipidomics data suggesting positive correlations between increase in DGDG and stromule formation were strengthened further upon observing increased stromules under phosphate starvation stress (Fig. S3) that is known to increase DGDG considerably in plant cells (Härtel et al., 2000; Andersson et al., 2003; 2005; Jouhet et al., 2004; Tjellström et al., 2008). Increased stromules were also observed in the aerial parts of the Arabidopsis *phosphate1* (*pho1)* mutant (Poirier et al., 1991; Fig.S3-C) shown to have nearly 170% more DGDG compared to wild type (Härtel et al. 2001). Another mutant, *hps1*(*hypersensitive to phosphate starvation1*) maintains a high level of sucrose internally due to overexpression of *SUCROSE TRANSPORTER2* (*SUC2*) and shows high sensitivity to Pi and N stress (Lei et al., 2011; Lei and Liu, 2011). Manipulating soluble carbohydrate accumulation in the *hps1* mutant by altering the concentration of sucrose in the growth medium was found to positively correlate with high stromule frequency in comparison to tpFNR-EGFP control plants (Fig. S3-D). High temperatures present yet another condition known to increase DGDG content in plants (Chen et al., 2006; Higashi and Saito, 2019). An increase in incidence of stromule formation was observed when tpFNR-EGFP plants were grown at 31°C instead of the usual 21°C temperature (Table 1). The strongest indication for DGDG involvement came from an absence of stromules in the *dgd1dgd2* double mutant with low DGDG content (Kelly et al. 2003) when they were treated with 40 mM sucrose in (Fig. 1H and Fig. S4-A-E). Notably, chloroplasts in guard cells that usually exhibit quite stable stromules in wild type plants did not extend stromules in the *dgd1dgd2* double mutant. Significantly, in addition to the blebbing chloroplast envelope and an apparent inability to extend stromules, the cells in *dgd1dgd2* double mutant showed many stroma-filled vesicles in the cytoplasm. The presence of stroma filled vesicles in the cytoplasm is a feature associated with senescent plastids (Costa et al., 2013; Ishida et al., 2014) and thus our observations suggested that reduced DGDG content weakens the plastid envelope. Additional evidence suggesting links between DGDG formation and stromules also came from a double mutant for *PHOSPHATIDIC ACID PHOSPHOHYDROLASE1 (PAH1)* and *PAH2* that is impaired in the eukaryotic pathway of galactolipid biosynthesis (Nakamura et al., 2009; Eastmond et al., 2010). By creating a double transgenic expressing tpFNR-EGFP and RFP targeted to the ER in the *pah1pah2* background we observed the characteristic expanded ER cisternae phenotype (Eastmond et al., 2010; Fig. S4-F, G). Upon treating these plants with 40 mM sucrose, while the chloroplasts exhibited an undulating form, long stromules were not observed Fig. S4G).

**Table 1:**
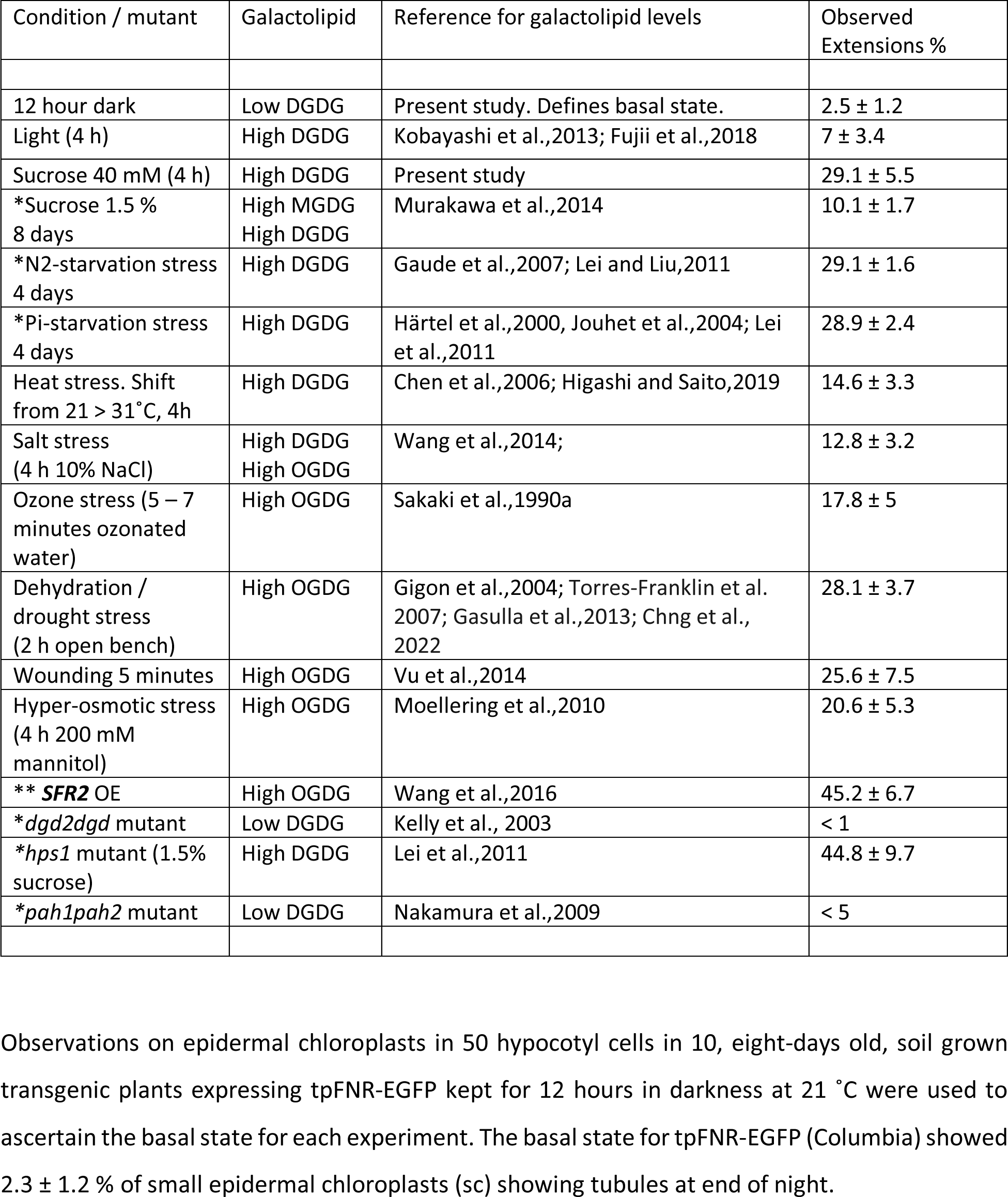
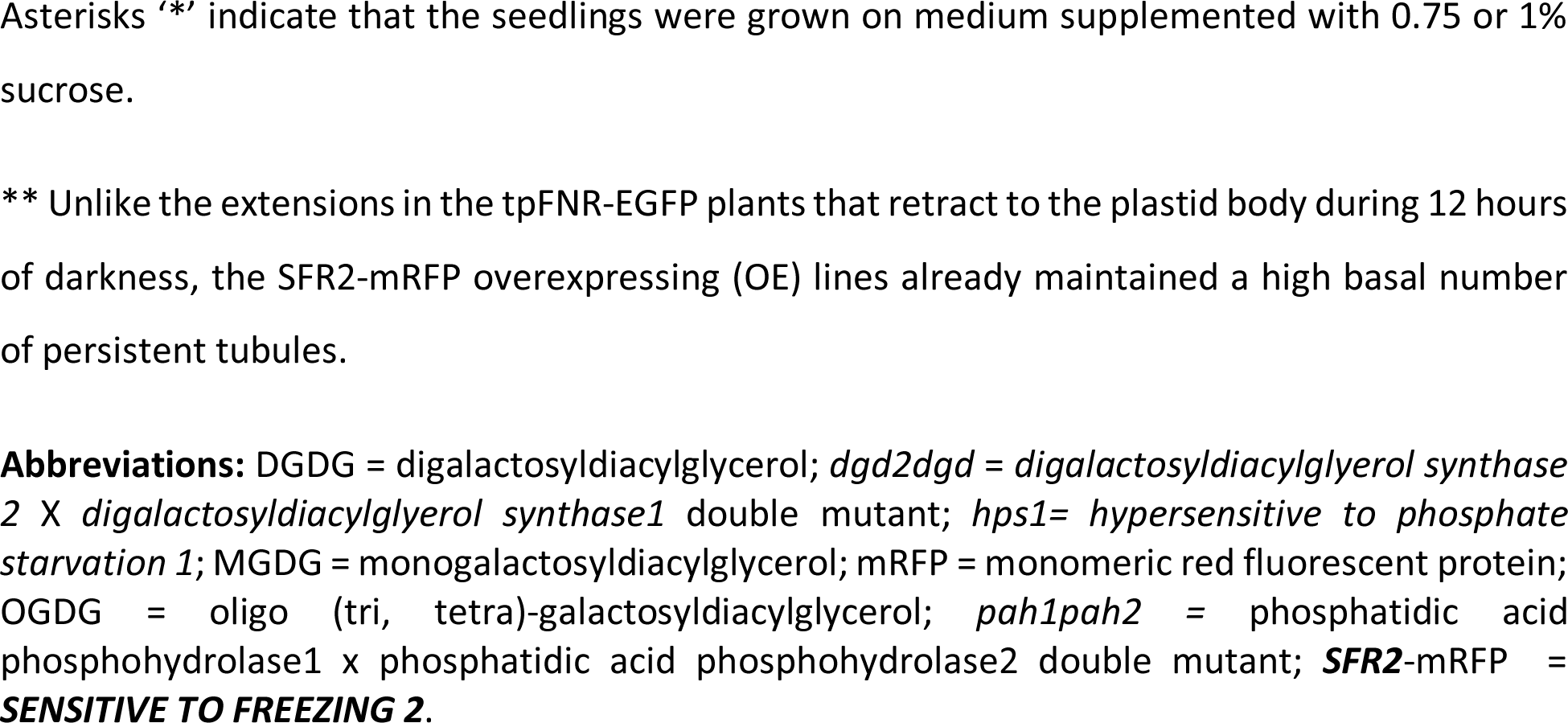
Conditions known to alter the level of bilayer-forming galactolipids in plant cells affect the formation of tubular extensions in plastids.

| Condition / mutant | Galactolipid | Reference for galactolipid levels | Observed Extensions % |
| --- | --- | --- | --- |
| 12 hour dark | Low DGDG | Present study. Defines basal state. | $2.5 \pm 1.2$ |
| Light (4 h) | High DGDG | Kobayashi et al.,2013; Fujii et al.,2018 | $7 \pm 3.4$ |
| Sucrose 40 mM (4 h) | High DGDG | Present study | $29.1 \pm 5.5$ |
| *Sucrose 1.5 %<br>8 days | High MGDG<br>High DGDG | Murakawa et al.,2014 | $10.1 \pm 1.7$ |
| *N2-starvation stress<br>4 days | High DGDG | Gaude et al.,2007; Lei and Liu,2011 | $29.1 \pm 1.6$ |
| *Pi-starvation stress<br>4 days | High DGDG | Härtel et al.,2000, Jouhet et al.,2004; Lei et al.,2011 | $28.9 \pm 2.4$ |
| Heat stress. Shift<br>from 21 > 31 °C, 4h | High DGDG | Chen et al.,2006; Higashi and Saito,2019 | $14.6 \pm 3.3$ |
| Salt stress<br>(4 h 10% NaCl) | High DGDG<br>High OGDG | Wang et al.,2014; | $12.8 \pm 3.2$ |
| Ozone stress (5 – 7<br>minutes ozonated<br>water) | High OGDG | Sakaki et al.,1990a | $17.8 \pm 5$ |
| Dehydration /<br>drought stress<br>(2 h open bench) | High OGDG | Gigon et al.,2004; Torres-Franklin et al.<br>2007; Gasulla et al.,2013; Chng et al.,<br>2022 | $28.1 \pm 3.7$ |
| Wounding 5 minutes | High OGDG | Vu et al.,2014 | $25.6 \pm 7.5$ |
| Hyper-osmotic stress<br>(4 h 200 mM<br>mannitol) | High OGDG | Moellering et al.,2010 | $20.6 \pm 5.3$ |
| ** *SFR2* OE | High OGDG | Wang et al.,2016 | $45.2 \pm 6.7$ |
| * *dgd2dgd* mutant | Low DGDG | Kelly et al., 2003 | < 1 |
| * *hps1* mutant (1.5%<br>sucrose) | High DGDG | Lei et al.,2011 | $44.8 \pm 9.7$ |
| * *pah1pah2* mutant | Low DGDG | Nakamura et al.,2009 | < 5 |

Observations on epidermal chloroplasts in 50 hypocotyl cells in 10, eight-days old, soil grown transgenic plants expressing tpFNR-EGFP kept for 12 hours in darkness at 21 °C were used to ascertain the basal state for each experiment. The basal state for tpFNR-EGFP (Columbia) showed $2.3 \pm 1.2$ % of small epidermal chloroplasts (sc) showing tubules at end of night.
Asterisks '\*\*' indicate that the seedlings were grown on medium supplemented with 0.75 or 1% sucrose.
\*\* Unlike the extensions in the tpFNR-EGFP plants that retract to the plastid body during 12 hours of darkness, the SFR2-mRFP overexpressing (OE) lines already maintained a high basal number of persistent tubules.
**Abbreviations:** DGDG = digalactosyldiacylglycerol; *dgd2dgd* = *digalactosyldiacylglycerol synthase* *2 X digalactosyldiacylglycerol synthase1* double mutant; *hps1*= *hypersensitive to phosphate* *starvation 1*; MGDG = monogalactosyldiacylglycerol; mRFP = monomeric red fluorescent protein; OGDG = oligo (tri, tetra)-galactosyldiacylglycerol; *pah1pah2* = phosphatidic acid phosphohydrolase1 x phosphatidic acid phosphohydrolase2 double mutant; **SFR2**-mRFP = **SENSITIVE TO FREEZING 2**.

Based on the strong correlations between increase in stromule numbers, ranging from five fold to nearly 50 fold, under several different conditions and mutants known to have high DGDG (Table 1; Fig. 3, 4) we concluded that there is a link between expansion of the OEM bilayer due to DGDG insertion and the extension of tubules. We reasoned that since the presence of DGDG with two galactose residues in the head group causes OEM extension, then a further increase in lipid headgroup size may cause the bilayer to stabilize and expand even more. Oligogalactolipids (OGDG) such as tri- and tetra-galactosyl diacylglycerol have larger headgroups than DGDG and add to the OEM-bilayer to increase its stability and structural integrity (Moellering et al., 2010; Roston et al., 2014). The effect of increasing OGDG on tubule extension was investigated next.

### SFR2-mRFP fusion reveals the OEM lying outside stromules

Our qPCR-based analysis showed that expression of SFR2, the gene responsible for GGGT activity leading to OGDG synthesis (Moellering et al., 2010), is substantially reduced following a 40 mM sucrose treatment (Fig. S5). Since OGDG production is triggered only under harsh stress (Sakaki et al., 1990a, 1990b; Moellering et al., 2010; Vu et al., 2015; Gasulla et al., 2019) an over-expression based approach was taken. Twelve stable transgenic lines expressing different SFR2 fluorescent fusion proteins (SFR2-mRFP, SFR2-EGFP and SFR2-mEosFP) under a constitutive CaMV35S promoter were created. In concurrence with earlier reports (Fourrier et al., 2008; Ferro et al., 2010) a clear highlighting of the plastid OEM was observed for plastids in all the lines. The SFR2-mRFP line #14 was identified as having the highest level of SFR2 expression (Fig. S5). Lipidomics analysis of this line revealed it having a nearly 17-fold increase in tri-galactosyldiacylglycerol (TGDG) compared to the wild type plants, along with significant increases in 34:2, 34:5 and 34:6 DGDG species (Fig. S6). The line thus had an overall relatively high ratio of bilayer-forming (DGDG +TGDG) lipids to the non-bilayer forming MGDG (Fig. 1I; Fig. S6).

A striking feature of the SFR2-mRFP transgenic overexpressing (OE) line #14 was the presence of very long tubules, ranging from 10 to 50 µm, extended by more than 65 % of sc, nearly 8 fold more than that normally observed for the tpFNR-EGFP plants (Fig. 1J and Fig. S5). Similar to the tpFNR-EGFP line the lc in SFR2-mRFP line, despite its strong, constitutive expression due to the CaMV35S promoter, retained a regular lens-like outline and did not form ectopic tubules (Fig. 1J). However, like the tpFNR-EGFP plants, the SFR2-mRFP OE line responded to 40 mM sucrose treatment by extending tubules from both sc and lc. This indicated that rather than SFR2 activity it was the 40 mM sucrose-induced increase in DGDG that increased the length of an existing tubule in these plants. Notably, in contrast to the near absence of tubules observed in tpFNR-EGFP plants after 12 hour darkness the tubules did not disappear in SFR2-mRFP OE plants (Fig.S5).

The creation of SFR2-mRFP plants the plastid OEM highlighted specifically allowed us to observe the membrane lying outside the IEM-delimited stromule. Considering that the IEM and the OEM differ considerably in their lipid compositions (Block et al., 2007), our working hypothesis was that conditions favouring addition of bilayer-forming lipids such as DGDG specifically to the OEM will affect the spatial relationship between the two membranes.

Therefore several double transgenic lines (Table S2) expressing differently coloured fluorescent proteins targeted to the stroma and to the OEM were created for their simultaneous visualization. Using separate lasers to distinguish the red and green fluorophores the IEM-delimited stromule and the OEM lying outside it could be clearly distinguished in lines expressing OEM-targeted SFR2-mRFP with stroma-targeted tpFNR-EGFP, an OEM-localized N-CHUP-EGFP (Oikawa et al., 2003) combined with stroma-localized CAC1-mRFP (Ke et al., 1997), N-CHUP-EGFP with stroma-targeted YFP (ptyk; Nelson et al., 2007), and OEM localized OEP7-GFP (Schleiff et al., 2001; Lee et al., 2001) combined with CAC1-mRFP (Table S2).

Upon assessing hypocotyl cells of 8 to 10 days old plants of the different double transgenic lines we observed a varying number of tubules where the stromule did not extend all the way to the tip and thus created an overhanging OEM-only region (Fig. 2A (control) versus 2B). Time-lapse imaging showed us that the spatial relationship between the stromule and its encasing OEM-tubule changed continuously. Focussing on the (N-CHUP-EGFP + CAC1-mRFP) and the (SFR2-mRFP + tpFNR-EGFP) combinations revealed that the frequency of plastids with an extra OEM-only portion (Fig. 2C) increased upon treatment with 40 mM sucrose for 4 hours in the dark. Based on our earlier findings (Fig. 1) these observations strongly suggested that the extra OEM-only extension upon sucrose treatment resulted from an increase in bilayer-forming DGDG.

**Figure 2.**
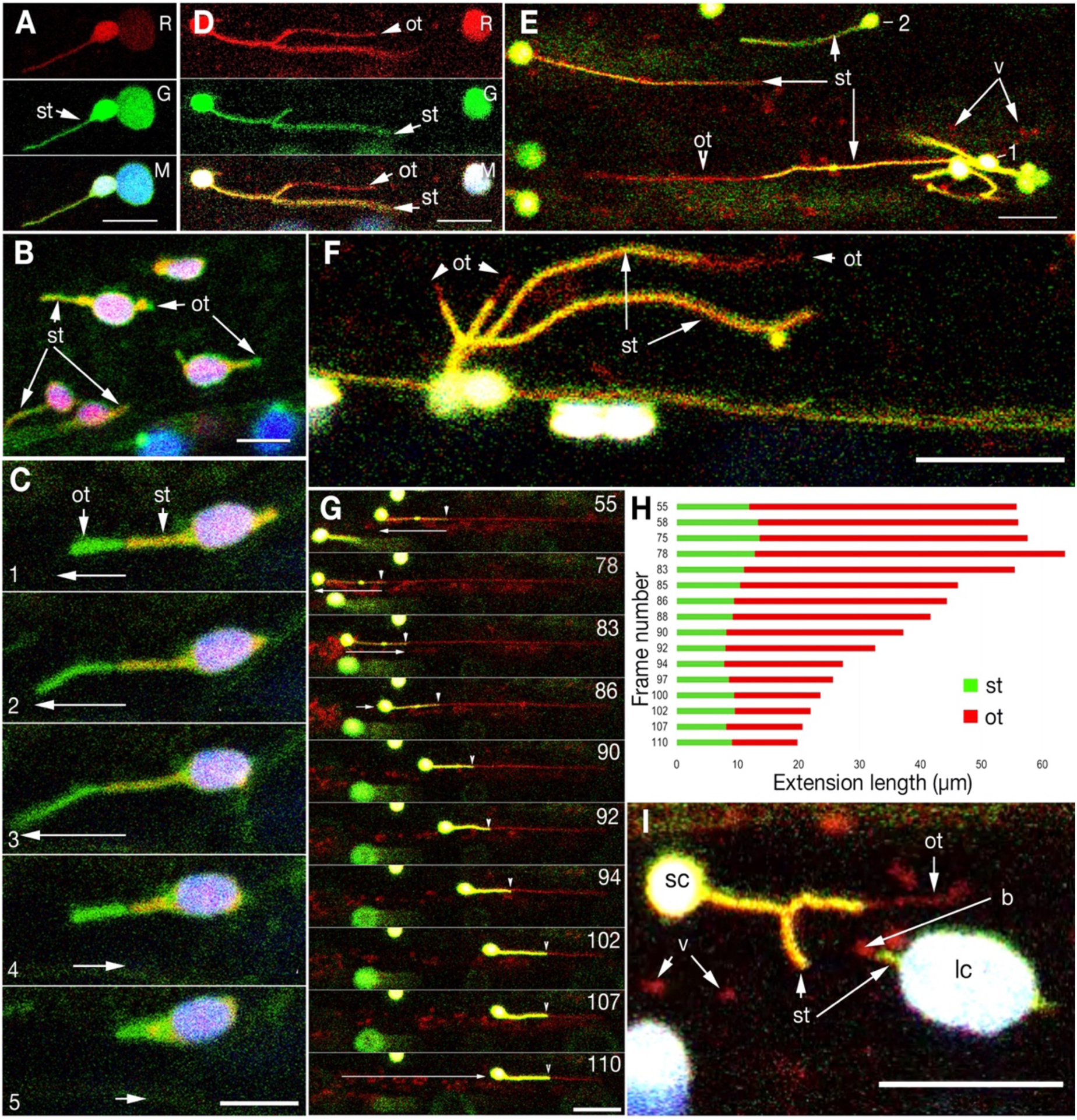
Double transgenic plants simultaneously expressing stroma and OEM targeted fluorescent fusions proteins reveal the spatial relationship between a stromule and an outer encasing-tubule with a high proportion of bilayer-forming galactolipids. **A.** Representative image from a transgenic line expressing stroma-targeted tpFNR-EGFP shows a stromule (st). There is slight bleed-through of EGFP and chlorophyll in the plastid body (pb) in the red (R) channel. B. Representative image from a double transgenic plant expressing N-CHUP-EGFP that highlights the OEM with stroma-targeted CAC1B-mRFP shows the green OEM extending beyond the red stromule. Note that the size of OEM extensions varies between different plastids. C. Sequential images from a time-lapse (x,y,t) series from a double transgenic plant expressing N-CHUP-EGFP + CAC1B-mRFP shows the tandem dynamic behavior of the stromule and extended OEM. D. Representative image from double transgenic line expressing stroma-targeted tpFNR-EGFP + OEM-targeted SFR2-mRFP shows the red fluorescent OEM tubule (ot) extending beyond the IEM delimited (arrowhead) stromule (st). E. A representative snapshot showing that at a particular time while some plastids (e.g. 1) may exhibit a long OEM extension (ot) at the tip of a stromule (st) another plastid (e.g. 2) may not exhibit the phenomenon. The presence of red vesicles (v) suggests that the OEM might be shed off into the cytoplasm. F. Snapshot showing the difference in size of OEM tubules (ot) extended beyond the stromules (st) by plastids in a cluster (Supplementary movie 4 shows the underlying dynamic behavior). G. Key sequential frames from an x,y,t time-lapse series (Supplementary movie 6) that shows the extension and retraction of a stromule within an OEM tubule stabilized by bilayer-forming DGDG and OGDG. Arrowheads indicate the limit to which the stromule extends within OEM tubule. Frames 55 to 78 depict stromule retraction while frames 83 onwards show stromule extension within the OEM tubule that progressively becomes shorter as red fluorescent OEM vesicles are shed into the cytoplasm. H. A kymograph depicting the changing spatial relationship between the stroma-filled, green fluorescent portion and the extended OEM (red) during the frames 55 to 110 shown in ‘G’ and in time-lapse supplementary movie 6. I. Snapshot from time-lapse series (supplementary movie 5) showing that treatment with 40 mM sucrose in the dark for 4 hours results in both small chloroplasts (sc) and large chloroplasts (lc) producing OEM protuberances (ot) and / or small blebs (b) of varying sizes that break off to form cytoplasmic vesicles (v). Size bars – B, C=5; A, D, E, F, G, I =10 µm.

By carefully charting the behaviour of the SFR2-mRFP #14 and the tpFNR-EGFP parent lines for the double transgenic combination we had established that the green fluorescent tpFNR-EGFP-highlighted stromule retracts towards the plastid body during 12 hours of darkness (Fig. 1C). By contrast the very long tubules extended by sc in the SFR2-mRFP OE line had their OEM highlighted and often persisted after 12 hours of darkness (Fig. 1J and Fig. S5). Reasoning that the SFR2-mRFP + tpFNR-EGFP double transgenic line would allow us to observe the dynamic behavior of a green fluorescent stromule inside a red fluorescent OEM tubule we observed hypocotyl cells in soil grown 8 days old plants soon after a 12 hour dark period.

Long, red fluorescent OEM-only extensions with lengths ranging from a micrometre to tens of micrometres beyond the stromule were observed (Fig. 2D-G; Movie S2, 3,4,5). Fifteen time-lapse recordings established that a stromule extends and retracts within an OEM-tubule (Fig. 2G, H; Movie S6). A treatment with 40 mM sucrose resulted in both sc and lc producing OEM blebs (e.g. ‘b’ in Fig. 2 I) as well as tubules. In addition, cells in the double transgenic lines showed an increase in red fluorescent vesicles in the cytoplasm (Fig. 2I ‘v’; Fig.S7, S8). The increase in vesicles was investigated next.

### Increased stress leads to the extra OEM being shed

While double transgenic SFR2-mRFP + tpFNR-EGFP plants taken from the dark showed OEM-only extensions of varying length within two hours of exposure to 110 ± 10 μmol m^-2^ s^-1^ photosynthetically active radiation (PAR) light a majority of the tubules in these plants reached lengths similar to those recorded for stromules in tpFNR-EGFP plants (Schattat and Klösgen, 2011; Barton et al., 2018). We reasoned that either these chloroplasts did not accumulate extra OEM or that the OEM that formed was lost over time. Starting from plants kept in the dark for 12 hours and exposing them to light at 15 minutes increments showed two non-exclusive mechanisms involved in the loss of OEM that had extended beyond the stromule. Accordingly, a stroma-filled region could advance towards the tip of an OEM-tubule and fill it up completely (Fig. 2G, H; Movie S6, S7). Alternately, the extra OEM could fragment at random locations along its length to produce small tubules and vesicles that were carried away into the cytoplasmic stream. The light-induced shedding became minimal once the extra OEM had been trimmed down to the region occupied by the stromule, suggesting that the presence of both IEM and OEM reinforces a tubule and limits OEM shedding.

Although a majority of lc did not exhibit long stromules, exposing plants to light continuously showed extensive blebbing of their OEM and sporadic budding of OEM vesicles (Fig.2I). A stress treatment that did not involve high light intensity was created by treating the double transgenic plants maintained under low light of 45 ± 10 μmol m^-2^ s^-1^ with 10% NaCl solution in water. OEM shedding increased within 5 minutes of the salt treatment.

The cytoplasmic vesicles lacked stromal contents and were exclusively red when observed in SFR2mRFP + tpFNR-EGFP plants or green in the case of N-CHUP-EGFP + CAC1mRFP double transgenics. This suggested their exclusive derivation from the OEM and set them apart from vesicles observed in cells with damaged or senescent plastids.

### OEM-shedding occurs in an ER-facilitated manner

Earlier investigations have established that the dynamic behaviour of stromules reflects their alignment with the cytoskeleton-dependent continuous reorganization of contiguous ER membranes (Schattat et al., 2011a; 2011b; Mathur et al., 2022; 2023). A double transgenic line expressing OEM-targeted SFR2-mRFP and GFP-targeted to the ER routinely showed OEM blebs in proximity to ER tubules (Fig. 3A). A triple transgenic line expressing stroma-targeted tpFNR-EGFP + OEM targeted SFR2-mRFP and YFP targeted to the ER lumen showed entire tubules (comprising of stromule and extended OEM tubule) enmeshed in and maintaining dynamic behaviour in tandem with the neighbouring ER (Fig. 3B; Movie S10). It was also observed that the dynamic tubules resulted from an OEM bleb being pulled out through its close alignment with contiguous ER (Fig. 3C Movie S8). Sporadically, the OEM tubules broke off and formed red fluorescent OEM-derived vesicles that became enmeshed independently in the ER (Fig. 3D) and moved in tandem with it into the cytoplasm (Movie S9). Further evidence supporting the formation of OEM vesicles came from transmission electron micrographs that showed separation of IEM and OEM in the plastid envelope with the possible folding of the detached OEM to form a vesicle (Fig. 3E). Since the OEM did not break at fixed intervals, long tubular OEM fragments that displayed dynamic, polygonal shapes were also observed in addition to the vesicles (Fig.S7-D, E). The dynamic behaviour of thee free OEM-derived fragments was attributed to their alignment with ER tubules.

**Figure 3.**
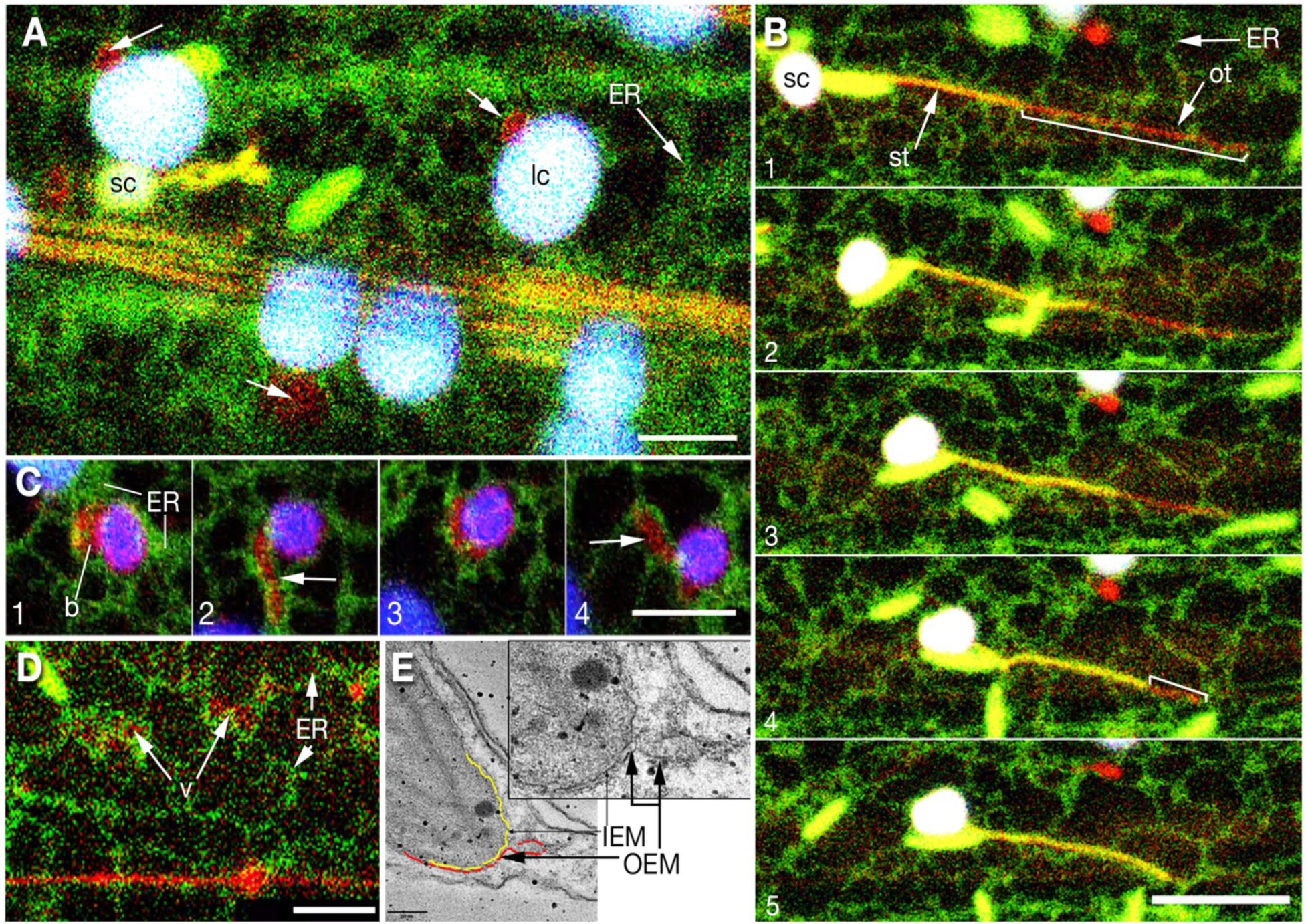
Tubular plastid extensions and OEM-derived vesicles are surrounded by a dynamic ER mesh. A. Representative image from a double transgenic line expressing SFR2-mRFP and ER lumen-targeted GFP shows the juxtaposition between OEM blebs in large chloroplasts (arrows) and ER tubules (arrow). B. Sequential frames from a time-lapse (xyt) sequence shows the shortening of an OEM tubule (ot), (extended OEM indicated by the length of the white line in panels 1 and 5), as a stromule (st) extends within it. Note that the entire tubule (stromule plus extended OEM) is enmeshed in the ER (green); Supplementary movie 10. C. Four sequential frames from an xyt time-lapse series (Supplementary movie 8) shows the extension of an OEM-bleb (b) to form a tubule (arrows in panels 2 and 4) through close association with contiguous dynamic ER. D. Representative image showing red fluorescent OEM-derived vesicles (v) free of the plastid and independently enmeshed in the ER (green; arrows) (Supplementary movie 9). E. A TEM image with IEM and OEM indicated in yellow and red, respectively (and inset) showing the opening out of the OEM and its refolding to possibly form an OEM-derived vesicle. Abbreviations: b – OEM-bleb; ER – endoplasmic reticulum; IEM – inner envelope membrane; lc-large sub-epidermal chloroplast; OEM – outer envelope membrane; sc-small epidermal chloroplast; v – vesicles. Size bars -E = 200 nm; A,C, D =5µm; B-10 µm

### A general strengthening of cellular membranes follows increase in OEM-localized galactolipids

A strong differentially permeable lipid bilayer is required to maintain organellar and cellular integrity. As assessed by transmission electron micrographs the SFR2-mRFP OE line #14 with a high content of bilayer-forming galactolipids had increased plastid OEM thickness (Fig. 5A, B, C). In addition, electrolyte leakage through membranes is widely used as a negative hallmark response to measure stress-induced injury as well as stress tolerance (Warren et al., 1996; Demidchik et al., 2014; Hincha and Zuther, 2014). Whether the 40 mM sucrose induced increase in DGDG also reduced ion-leakage was assessed and exhibited significantly reduced ion-leakage (Fig. 5C, D). In comparison to the wild type plants the *dgd1* mutant with nearly 90% reduction in DGDG production (Dörmann et al., 1995; 1999) exhibited significantly increased ion leakage (Fig. 5E).

It was concluded that conditions that enhance bilayer-forming, plastid-derived galactolipids such as DGDG and OGDG in plant cells reinforce cellular membranes and reduce ion leakage.

## Discussion

Since their recognition as essential organelles of the viridiplantae (Schimper (1882) the dynamic behaviour of plastids has been a major area of investigation. The formation of long tubular, transient protuberances of the plastid envelope that appeared sporadically, extended into the cytoplasm and retracted towards the plastid body presented an specially intriguing conundrum (Wildman et al., 1962; Hongladarom et al., 1964). Targeting a GFP to the stroma revealed that the tubules were filled by the fluid stroma (Köhler et al., 1997). The stroma-filled tubules were called stromules (Köhler and Hanson, 2000).

Folowing their discovery the stromules have been fascinating subjects for myriad cell biological investigations (Gray et al., 2001; Natesan et al., 2005; Hanson and Conklin, 2020) but the aims of uncovering the mechanism underlying their formation and their function within the plant cell have remained elusive. In retrospect, two reasons for limiting our understanding can be identified; one, the sporadic and inconsistent nature of the phenomenon that made it difficult to subject it to a strong characterization, and, two, not giving sufficient recognition to the fact that that the stromule is actually delimited by the inner envelope membrane (IEM) and is encased within a robust outer envelope membrane (OEM). Notably, it is the OEM and not the stromule that is directly in contact with, and continuously interacts with the rest of the cytoplasm.

By recognizing the two limiting factors and resolving them our work provides fresh insights on the phenomenon of tubule formation in plastids.

The first step towards understanding the formation of tubules from plastids was define a condition where they cannot be observed and then to identify a second condition where they can be elicited at a high frequency. Extended stromules were nearly absent in chloroplasts in hypocotyl cells of soil-grown Arabidopsis tpFNR-EGFP transgenic line (Schattat et al., 2011a) after a 12-hour period of darkness. Tubules could be induced at a high frequency upon treating the plants with 40 mM sucrose in the dark for 3 to 4 hours (Barton et al., 2018). The consistent and easily reproducible method paved the way for comparative cell biological, molecular and lipidomic analyses.

The sucrose induced extension of tubules in the dark progressed from a tiny protuberance of the plastid envelope to a several micrometers long tubule within 4 hours. While underscoring the relatively short time required to elicit the phenomenon, the observations raised questions about the significance of many reports where the experiments span many more hours and sometimes extend over several days. Moreover, the observations also established that exposing chloroplasts to light is not a requirement for tubule extension. This also suggests that triggering of reactive oxygen species (ROS) by strong light, considered a strong inducer of stromules (Brunkard et al., 2015; Exposito-Rodriguez et al., 2017), may only be considered an augmenting factor. However, as shown by the sucrose treatment, an increase in cytosolic sucrose levels turned out to be a strong inducer of tubules (Fig. S1). The 40 mM sucrose concentration is physiologically relevant and lies within the estimated sucrose levels reached in photosynthetically active plant cells (Winter et al., 1994; Gerhardt et al., 1987). The response to sucrose also explained why plastids in non-photosynthetic cells have been observed extending stromules readily (Natesan et al., 2005; Barton et al., 2018; Hanson and Conklin, 2020). A rise in cellular sugar is also seen to accompany many other abiotic stresses including drought, salinity, oxidative and temperature stresses (Baxter et al., 2006; Wingler and Roitsch, 2008; Rosa et al., 2009) that have also been shown to result in the extensions. Sugar release and accumulation also occurs in response to pathogens, and endogenous or exogenous sugars can trigger many pathways associated with pathogen resistance and biotic stresses (Herbers et al., 1996; Thibaud et al., 2004; Gómez-Ariza et al., 2007). Most importantly, tubule induction by sucrose as well as glucose (Schattat and Klosgen, 2011) suggested a heterotrophic phenomenon.

One of the effects of sucrose on biological membranes is to stabilize them through interactions with membrane lipids (Moiset et al., 2014). However, at relatively low concentrations such as used in our experiments sugars can also cause membranes to expand and become thinner by insertion between the lipid heads, and thereby reduce membrane bending rigidity (Andersen et al., 2011; Dimova, 2014). We reasoned that our observations of sugar-induced stromule extension could be pointing to changes in the properties of plastid envelope membranes that are largely defined by the characteristic galactolipid compositions of the IEM and OEM (Dorne et al., 1982; 1985;Block et al., 2007). Our analyses strongly suggested that DGDG, a bilayer-fostering galactolipid increases following treatment with sucrose (Fig. 1; Fig.S2). Interestingly, a change in sucrose concentration also alters phosphate levels in a cell (Karthikeyan et al., 2007; Lei et al., 2011). Phosphate starvation stress specifically, has been suggested to stimulate an increase in DGDG production to compensate for loss of membrane phospholipids (Essigmann et al., 1998; Härtel et al.,2000; 2001; Kelly and Dormann, 2002; Jouhet et al., 2004). Indeed, our observations showed that plastid tubules increased under Pi-starvation stress. Other conditions including increase in temperature, and dehydration that are known to augment DGDG levels in plants also increased the incidence of tubule extensions (Table1; Fig. S3). Strong evidence supporting a role for DGDG in stromule extension when *dgd1dgd2* double mutant plants shown to have greatly reduced levels of DGDG (Kelly et al., 2003), were treated with 40 mM sucrose but failed to show tubules (Fig. 1 and Fig. S4).

Following the positive correlation between OEM strengthening and DGDG formation on the plastid OEM upon sucrose treatment, the effect of a further increase in OEM stability was tested. Oligogalactolipids (OGDG) as tri- and tetra-galactolsyldiacylglycerol with larger headgroups than DGDG have been shown to increase membrane-stability but their production is triggered naturally only under harsh stress (Benning and Ohta; 2005; ; Vu et al., 2005; Gasulla et al., 2019; Barnes et al., 2016; 2023). We circumvented the requirement for harsh stress by overexpressing the SFR2 gene, responsible for GGGT activity leading to OGDG production (Moellering et al., 2010) under a strong constitutively active CaMV-35S promoter. Increased SFR2 expression levels and OGDG amounts (Fig. 1 and Fig. S5, S6) in the stable transgenic lines allowed us to avoid artefacts resulting from general membrane breakdown under harsh stress while being able to observe the effects of plastid-OEM strengthening.

A clear effect of increased OGDG was the increased length of the plastid tubules. Although the SFR2-mRFP labelled the OEM in all plastids, the formation of tubular extensions was mostly limited to small chloroplasts (sc) while the large sub-epidermal chloroplasts (lc) did not form tubules. The distinctly different tubule extension behaviour between sc and lc in stable transgenic SFR2-mRFP OE plants clearly argued against the transient overexpression-based idea which suggests that stromule-like tubules form due to over-expression of any OEM-localized protein (Breuers et al., 2011). Similar to the SFR2-mRFP OE lines, stable, non-injured transgenic plants expressing OEM-localized N-CHUP-EGFP and OEP7-GFP proteins also did not show excessive tubule formation from large chloroplasts. Nevertheless, in all the transgenic lines for OEM-localized proteins both sc and lc extended tubules when subjected to 40 mM sucrose treatment that has been shown to increase DGDG levels. It is noteworthy that most transient expression-based experiments are carried out using agroinfiltration of tobacco leaves followed by fluorescent microscopy-based observations on excised tissue. Considering the analysis reported by Vu et al.,(2014, 2015) wounding triggers an increase in SFR2 expression and the presence of OGDG in a cell. There is thus a possibility that tissue handling and wounding of the leaf tissue observed by Breuers et al., (2011) triggered OGDG production and resulted in an abnormal stability of tubule membranes that may have contributed to their conclusions.

A second observation on plants expressing OEM-targeted probes N-CHUP-GFP and SFR2-mRFP was that more than 50 % of the tubules extended from small chloroplasts (sc) were longer than stromules observed in plants expressing stroma-targeted tpFNR-EGFP (Fig. 1; Fig. S5). We realized that although tubules were observed in tpFNR-EGFP, SFR2-mRFP OE and N-CHUP-GFP lines, the fusion probes involved actually highlighted different sections of the double-membraned chloroplast envelope. A perusal of literature showed that while stroma-targeted fluorescent proteins have been used to correctly identify stromules, little consideration has been paid to the fact that the stromule is actually confined within the plastid IEM and is thus separated from the OEM by an inter membrane region (IMR; Fig. 1B; Fig. 5). While under normal conditions of growth and development the IEM and OEM may remain very close, in the range of 5 to 10 nm from each other (Keegstra et al., 1984; Cline et al., 1985; Cremers et al., 1988; Engel et al., 2015) in comparison to the IEM the OEM has a high DGDG content (Block et al., 2007; Benning 2009). As shown through our observations on double transgenic plants highlighting stromules and the OEM simultaneously conditions that overly favour addition of bilayer-forming DGDG to the OEM allow it to extend extend beyond the stromule (Fig 2; Fig. S5, S7). Moreover, the 12-hour dark period when the plastid body did not exhibit extended stromules (Fig. 1C), showed long OEM-only tubules without an inner stromule. Importantly, treatment with 40 mM sucrose (that specifically increases DGDG), resulted in OEM-tubule extension accompanied by an increase in OEM-derived vesicles in the cytoplasm. The presence of vesicles suggested that extra, DGDG-rich OEM could be shed and was confirmed using high light and salt stress (Fig. 3; Fig. S7E, F; Movies S8,S9,S10). It was concluded that despite being stabilized by the presence of bilayer-forming galactolipids, portions of the OEM that are not internally supported by an IEM are shed off into the general cytoplasm. Although the OEM shedding observed by us suggested similarity to the tip-shedding phenomenon described earlier for plastids using differential interference contrast (DIC) microscopy (Gunning, 2005), the OEM-derived vesicles did not have stromal contents. They thus differed from the small vesicles and senescence associated vacuoles (SAVs) containing stromal proteins that are normally observed in senescing leaves (Wada et al., 2009; Costa et al., 2013).

More importantly, the OEM-derived vesicles released into the cytoplasm, possibly by a cytoskeleton and ER mediated mechanism, closely associated with the endoplasmic reticulum (ER) membranes (Fig. 3; Movies S8, S9, S10). As part of the cytoplasm the galactolipid-rich vesicles could freely mingle with other cytoplasmic components and compartments. Indeed, there are earlier reports of general membrane reinforcement by galactolipids in response to different stresses including dehydration (Gigon et al., 2004; Moellering et al., 2010; Gassulla et al., 2013; Chng et al., 2022) and phosphate starvation stress (Härtel et al., 2000; Andersson et al., 2003,2004, 2005; Jouhet et al., 2004; Nakamura et al., 2009). Our ion-leakage based anlysis showed positive correlations between membrane reinforcement and reduction in ion leakage from plant cells (Fig. 4). While acknowledging that due to overexpression by a strong CaMV35S promoter the number and frequency of OEM-vesicles being released into the cytoplasm may be more than usual in our stable double transgenic plants, we point out that even at a very low rate of release, such as may occur diurnally under increased cellular sucrose levels, the fusion of galactolipid-rich, OEM-derived vesicles may serve to reinforce other organelle membranes.

**Figure 4.**
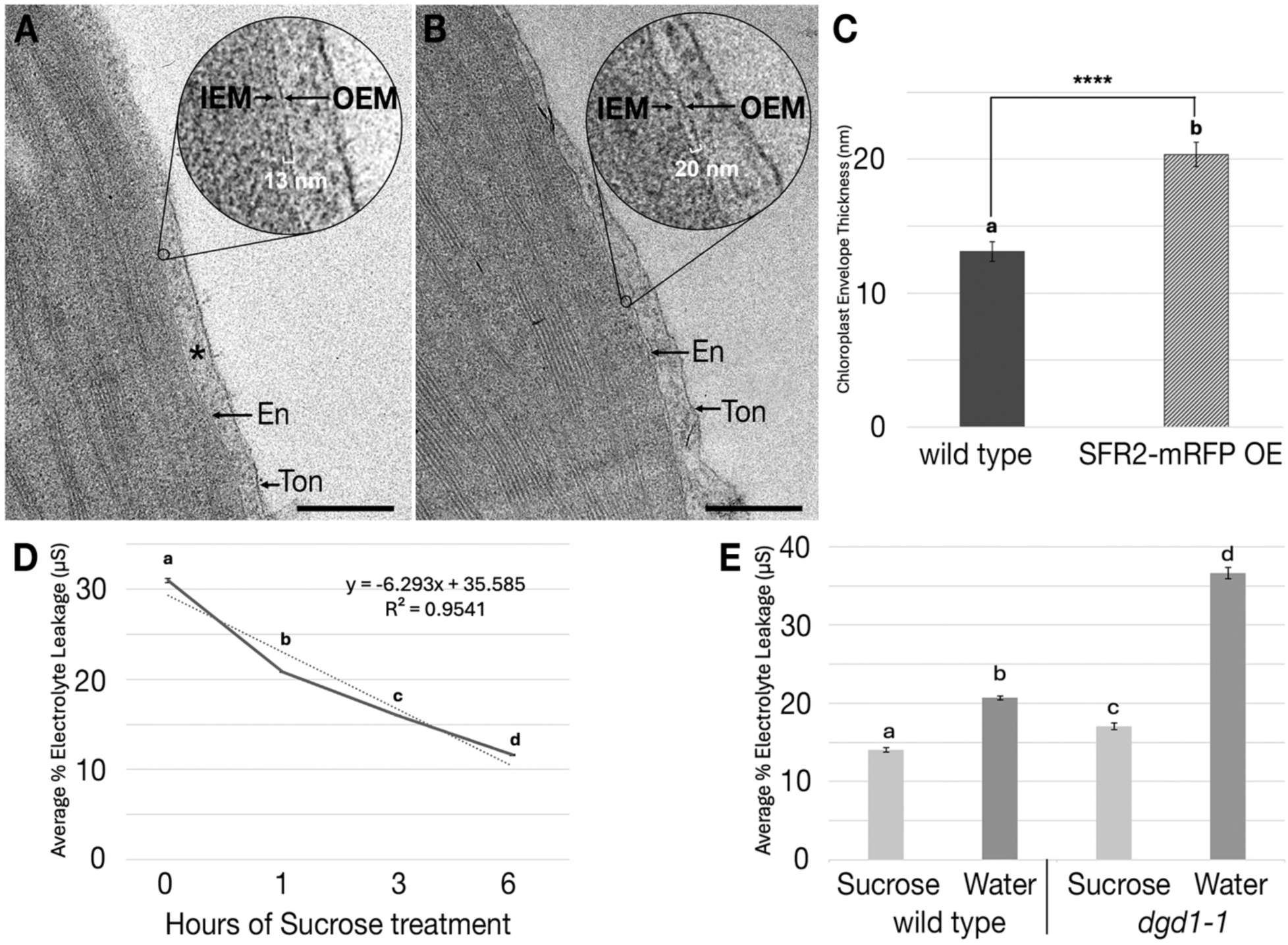
Transmission electron microscopy and electrolyte leakage measurements suggest that increase in bilayer-forming galactolipids reinforce cellular membranes. **A.** B. Transmission electron micrograph of wild type (ecotype Columbia) and B. SFR2-mRFP OE #14 mesophyll chloroplasts with emphasis on the envelope membrane. Asterisk = cytosol, arrows indicate chloroplast envelope (En), vacuolar tonoplast (Ton), inner envelope membrane (IEM), and outer envelope membrane (OEM). Scale bar = 400 nm. C. Bar graph depicting average chloroplast envelope thickness ± SE of 4 biological replicates, specifically the distance between IEM and OEM. The chloroplast envelope is thicker in the SFR2-mRFP OE line compared to wildtype. Each replicate measured all chloroplasts within 15 mesophyll cells, each chloroplast envelope was measured with 3 technical replicates each for a total of 180 measurements per plant line. Asterisks indicate significant differences within the treatment by a Student’s two-tailed t-test where **** is p < 0.0001. Varying letters indicate significant differences as determined with a Tukey’s HSD test at the p <0.01 level of significance. D. Sucrose treatment strengthens membranes and reduces electrolyte leakage suggested by percent electrolyte leakage (EL measured in µSiemens) between 7-day old *A. thaliana* seedlings which had been dark treated for 18 hours prior to water and 40 mM sucrose treatments. Low % EL indicates high membrane integrity with minimal ion leakage, higher % EL indicates weaker membrane integrity with high ion leakage. The 0-time point control placed seedlings directly into measurement milli-Q water. The treatments were: pre-treating plants with 40 mM sucrose for: 1 hr, 3 hours, and 6 hours before rinsing and transferring to the IEC measurement tubes. Error bars represent average electrolyte leakage ± standard deviation; n=12, varying letters represent Tukey’s HSD test where p<0.05. E. Bar graph showing that membranes in the *dgd1-1* mutant are significantly leakier than Columbia. Sucrose pre-treatment reduces membrane leakiness. Error bars represent average electrolyte leakage ± standard deviation; n=12, varying letters represent Tukey’s HSD test where p<0.01.

While our ongoing experiments are aimed at exploring the nature of interactions between plastid OEM-derived vesicles and other organelles and the possibility that specific membrane contact site fostering proteins might be involved, our present findings are summarized in Fig. 5. Accordingly it is suggested that the mechanism underlying tubule extension from a plastid involves a localized increase in bilayer-forming galactolipids such as DGDG on the plastid OEM. Formation of an OEM tubule creates a narrow conduit within which the IEM-delimited stroma flows to form a stromule. When the stromule retracts towards the plastid-body, portions of the OEM tubule that are no longer reinforced by the IEM can break off and be shed into the cytoplasm. The galactolipid rich vesicles freely associate with other cellular membranes. Since the tubules extend far into the cytoplasm, our observations strongly suggest that they have a role in dispersing plastid derived galactolipids in the cell. In turn, the association of galactolipids to existing cellular membranes could enhance membrane stability and serve to increase plant cell resilience to stress.

**Figure 5.**
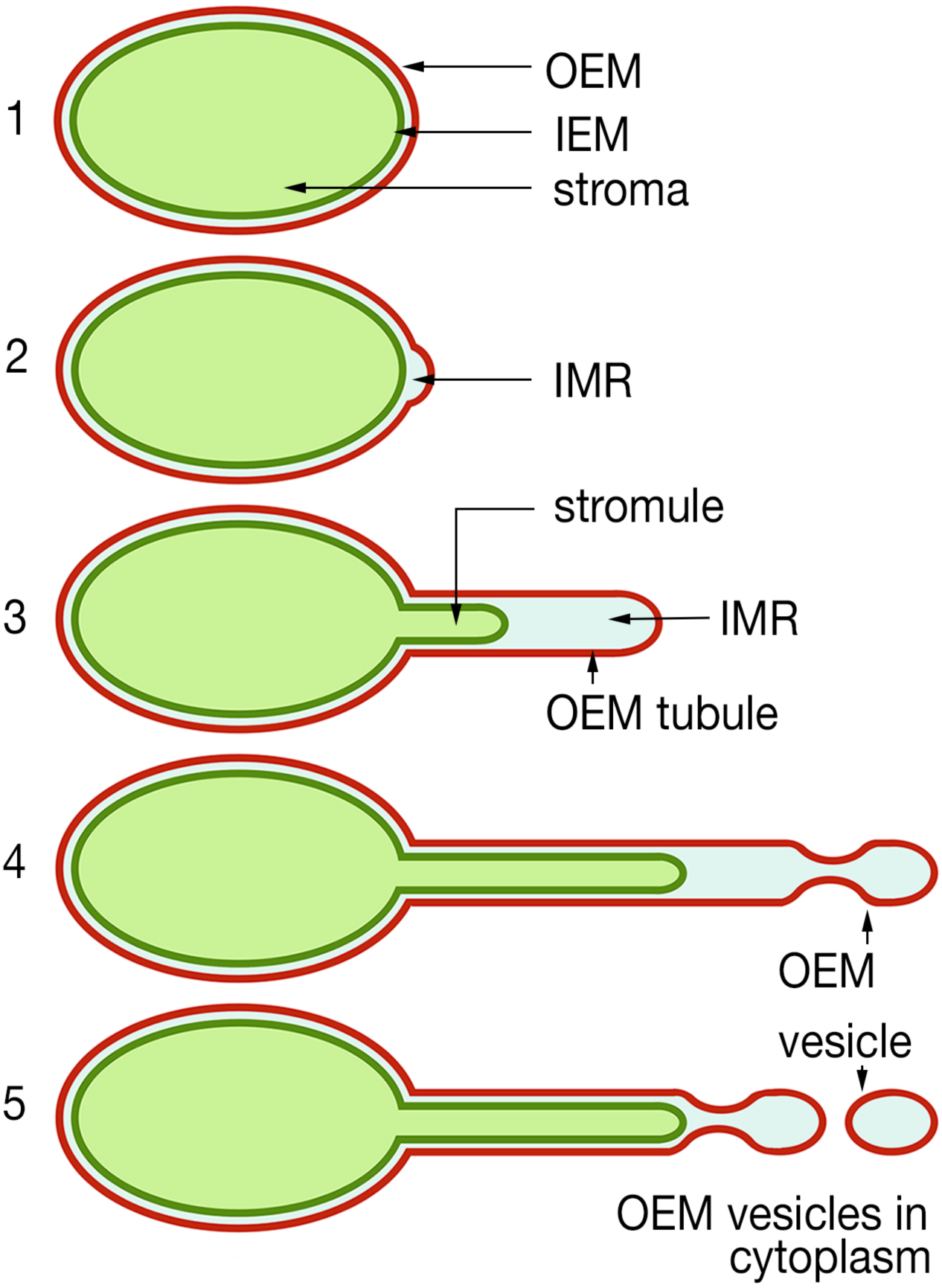
Diagrammatic depiction summarizing findings on tubular plastid extensions and release of bilayer-forming galactolipid rich vesicles into the cytoplasm. 1. A typical plastid in viridiplantae has a stroma-filled interior that is confined within the inner envelope membrane (IEM) which is itself encased within the outer envelope membrane (OEM). Compared to the IEM the OEM is rich in bilayer-forming galactolipids. 2. A thin intermembrane region (IMR) separates the IEM from the OEM. 3. The plastid envelope extends to form tubular protuberances under diverse stress. Targeting a fluorescent protein to the stroma highlights the IEM-delimited stroma and led to the stroma filled tubule being called a ‘stromule’. However, conditions that specifically favour addition of bilayer-forming lipids, such as DGDG, to the OEM cause the OEM-tubule to extend beyond the stromule. 4. The extended OEM-only portion of a tubule lacks an internal reinforcing membrane and is prone to pinching and breaking through cytoskeleton and ER mediation to form vesicles. 5. OEM-derived vesicles, rich in bilayer-forming galactolipids reach the cytoplasm where they may mingle with other cellular membranes.

## Supporting information

Single Supplementary file

## Author contributions

JM conceived, planned the experiments and carried out confocal imaging, LMJC carried out the TEM, created several SFR2-fusion constructs and genetic crosses and carried out experiments on mitochondria-plastid interactions, MB and LW analysed the lipidomics data, JC and KS carried out the PCR-based investigations, KB and MW carried out experiments on sugar, different nutrient stresses, NM created, characterized and maintained all transgenic lines and Arabidopsis mutants.

## Acknowledgments

We thank G. Wiedenmann for mEosFP, K. Oikawa for N-CHUP-EGFP, R. Mullen for OEP7-GFP, P. Eastmond for *pah1pah2* double mutant, D. Liu for *hps1* mutant, and acknowledge the help of Alyssa Clews in lipid extraction and TLC.

## Funding

The investigations from 2010 to 2025 were funded by Discovery grant RGPIN-2021-02448 to JM by the Natural Sciences and Engineering Research Council of Canada (NSERC). Instrument acquisition and lipidomics method development at KSU is supported by National Science Foundation (EPS 0236913, MCB 1413036, MCB 0920663, DBI 0521587, DBI1228622), Kansas Technology Enterprise Corporation, K-IDeA Networks of Biomedical Research Excellence (INBRE) of National Institute of Health (P20GM103418), and Kansas State University.

## Conflict of interest

The authors declare no competing interests.

## Data availability

All materials presented in this study are available upon request.

## Single Supplementary FILE

**Supplementary Table 1:**
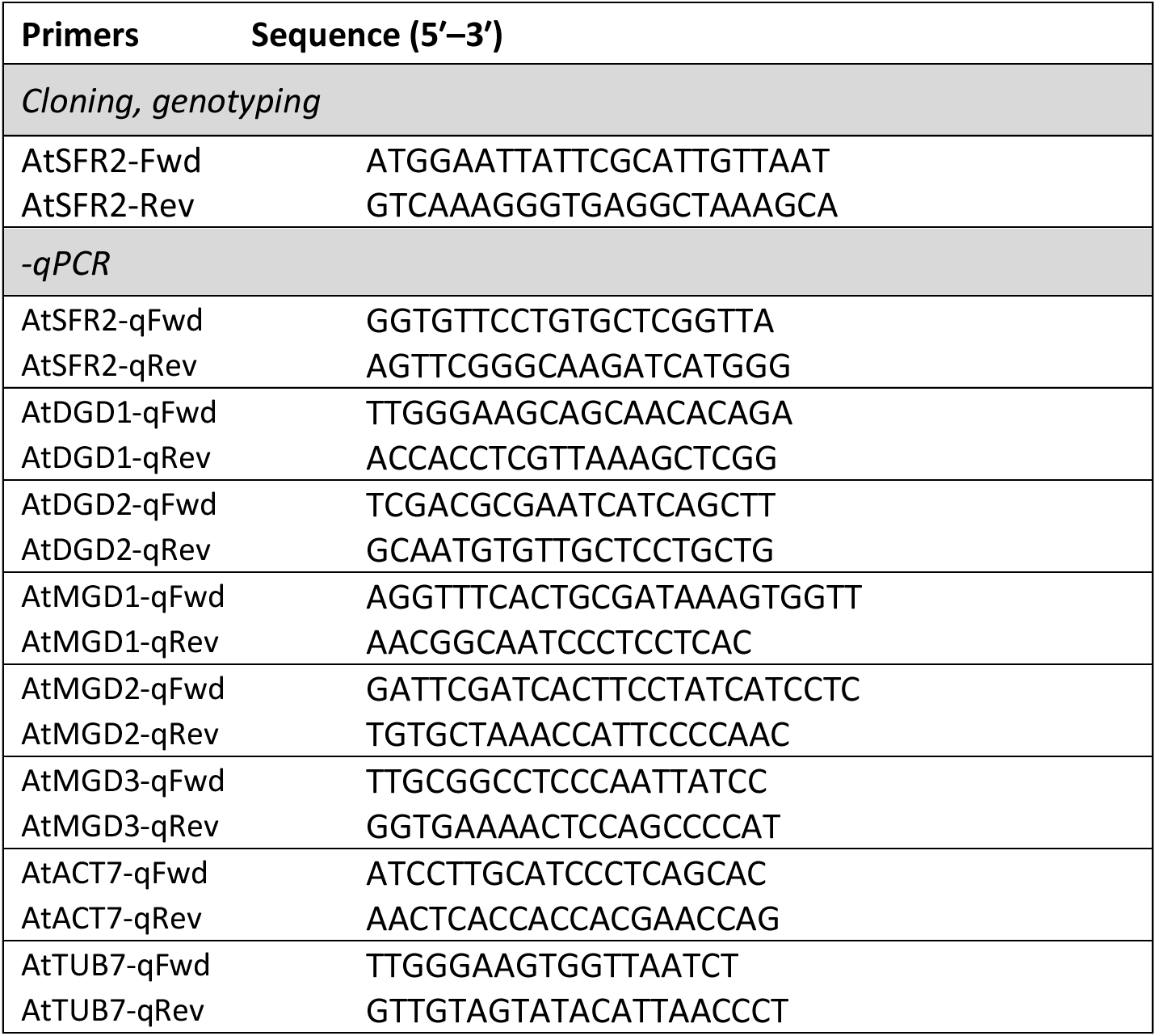
Gene-specific primers used for qPCR.

**Supplementary Table 2:**
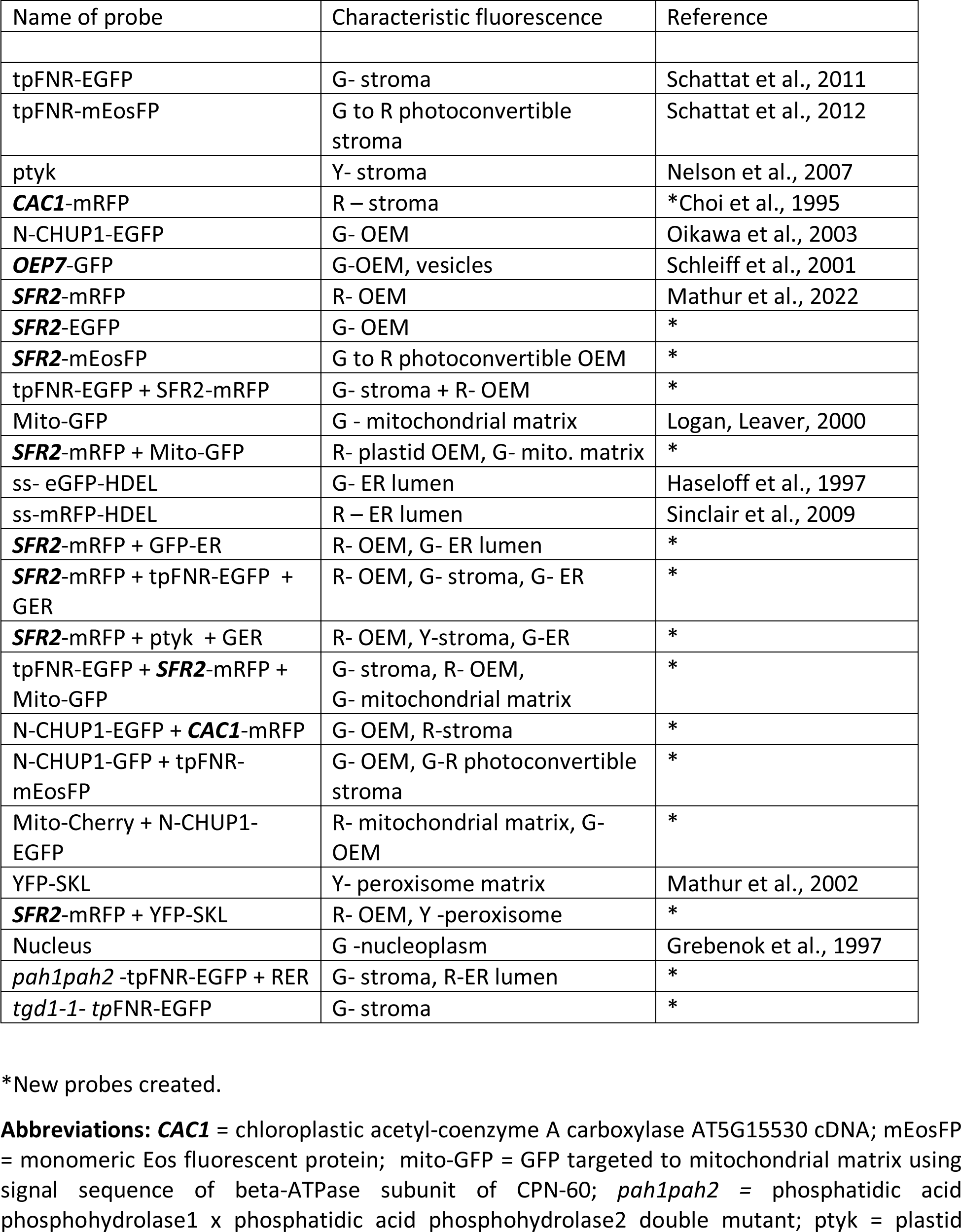
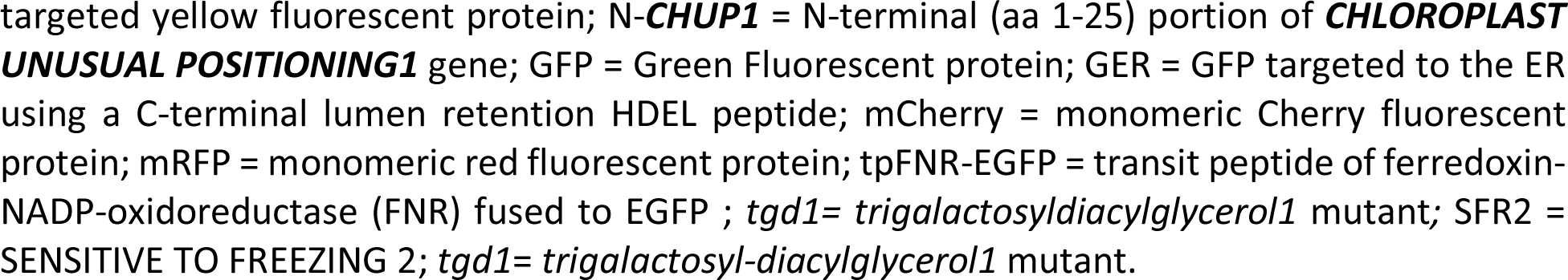
Probes used in the present study.

**Supplementary Figure 1.**
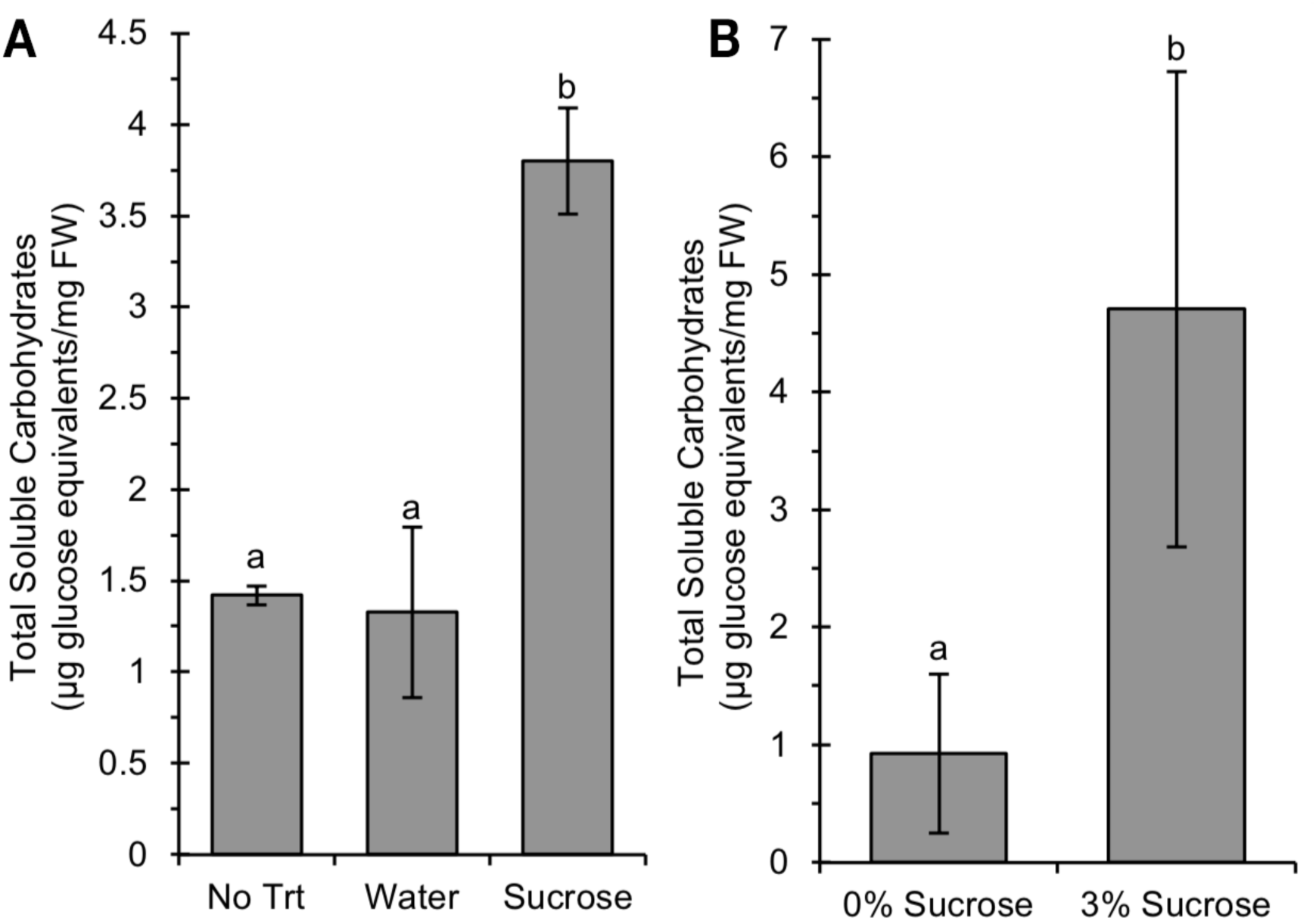
Estimation of total soluble carbohydrate levels. A. 3 h treatment with 40 mM sucrose resulted in higher soluble carbohydrate levels in the above-ground tissue than the water control and no treatment controls (both p<0.01, N=4). No difference was seen between the untreated and water treated plants (p=0.48, N=4). B. Plants grown on media containing 3% sucrose had higher total soluble carbohydrate levels than those grown on 0% sucrose media (p=0.009, N=3). Error bars indicate the 95% confidence interval of the mean. For carbohydrate analysis (N= x) represents the number of tissue samples, each containing the aerial portion of thirty seedlings.

**Supplementary Figure 2.**
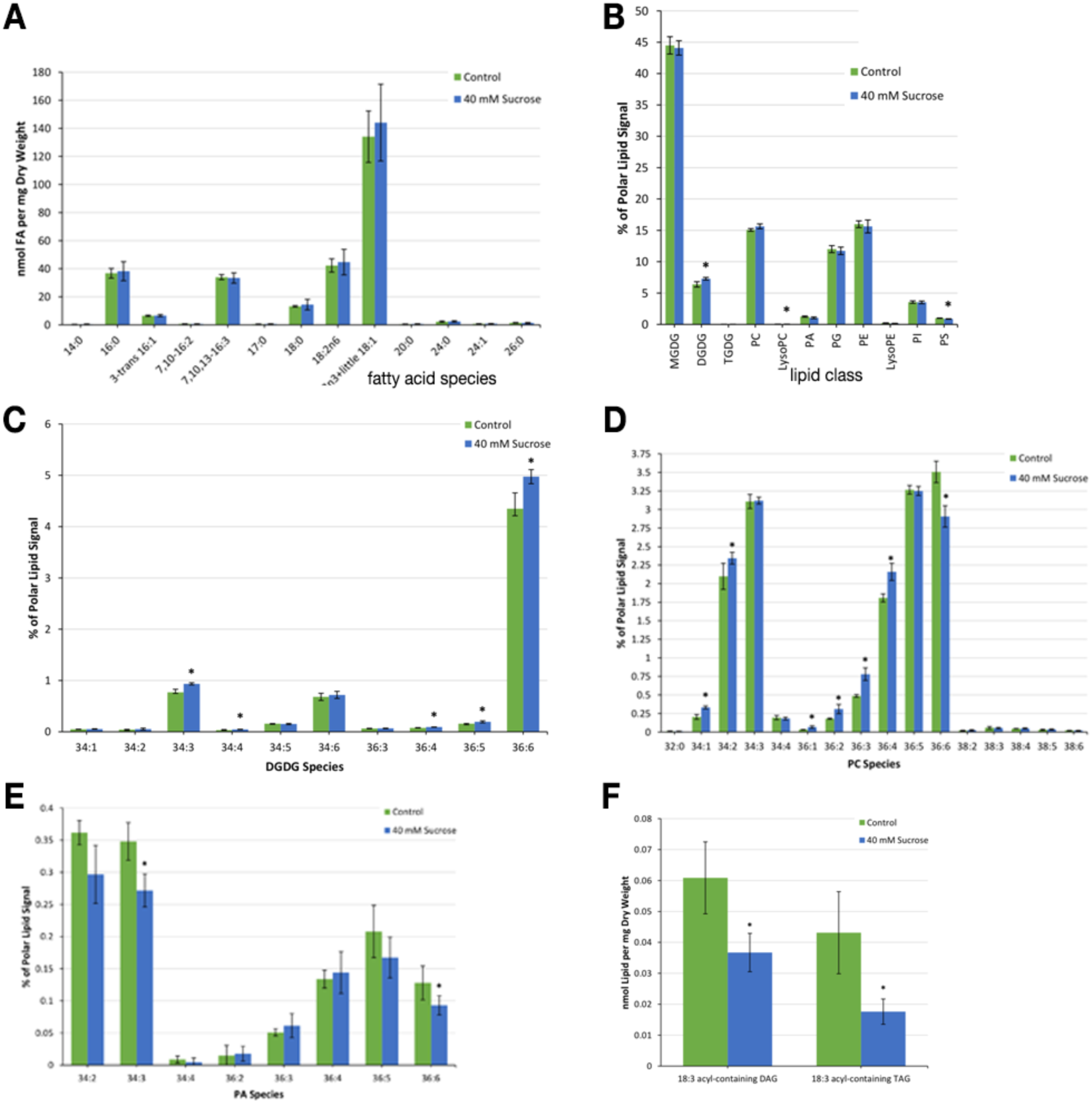
Fatty acid and lipid profiles following water and sucrose treatment of tpFNR-EGFP expressing transgenic plants (Columbia ecotype) **A.** Comparison of fatty acid profile (in nmol of FA detected per mg dry weight) for *A. thaliana* shows no significant changes following treatment with water and with 40 mM sucrose in the dark for 3 hours. Averages ± standard deviation calculated from five independent samples in 40 mM sucrose treatment and three independent samples in the control. Asterisks indicate statistically significant differences between treatments as determined by Student’s t-tests (P < 0.05). **B.** Polar lipid content in *A. thaliana* under a 40 mM sucrose treatment with respect to a water control. Following overnight darkness, rosette leaves from 31-day-old soil-grown plants were subjected to 3 hours of darkness in either water or 40 mM sucrose and lipids were subsequently extracted. Lipid profiles were quantified using ESI-MS, (Shiva et al., 2013) at the Kansas Lipidomics Research Center Analytical Laboratory. Averages ± standard deviation were calculated from five independent samples in 40 mM sucrose treatment and water controls. Asterisks indicate statistically significant differences between treatments as determined by Student’s t-tests (P < 0.05). Significant increase in DGDG. **C.** DGDG lipid species content in *A. thaliana* under a 40 mM sucrose treatment with respect to a water control in rosette leaves from 31-day-old soil-grown plants following overnight darkness. Leaves were subjected to 3 hours of darkness in either water or 40 mM sucrose before lipid extraction and quantification using ESI-MS. Averages ± standard deviation were calculated from five independent samples. Asterisks indicate statistically significant differences between treatments as determined by Student’s t-tests (P < 0.05). Significant increase in DGDG 34:3, 34:4, 36:4, 36:5 and 36:6. **D.** PC lipid species content quantified using ESI-MS for *A. thaliana* leaves treated with 40 mM sucrose treatment and a water control. Averages ± standard deviation based on 5 independent samples with asterisks indicating statistically significant differences between treatments as determined by Student’s t-tests (P < 0.05). While several PC species exhibited slight increases a significant decrease in PC 36:6 suggested it as a major source for DGDG 36:6. Although not shown here, 18:3 Lyso PC also decreased significantly under sucrose treatment compared to control. **E.** PA lipid species content in *A. thaliana* under a 40 mM sucrose treatment with respect to a water control. Asterisks indicate statistically significant differences between treatments as determined by Student’s t-tests (P < 0.05). Compared to controls, 34:3 and 36:6 PA decreased significantly following sucrose treatment. **F.** 18:3 acyl-containing non-polar lipid content in *A. thaliana* under a 40 mM sucrose treatment with respect to a water control (in nmol of lipid detected per mg of dry weight). Following overnight darkness, rosette leaves from 31-day-old soil-grown plants were subjected to 3 hours of darkness in either water or 40 mM sucrose and lipids were subsequently extracted. Lipid profiles were quantified using ESI-MS. Averages ± standard deviation calculated from five independent samples with asterisks indicating statistically significant differences between treatments as determined by Student’s t-tests (P < 0.05). Both 18:3 acyl-containing DAG and TAG decreased following sucrose treatment. Abbreviations: DAG- diacylglycerol; MGDG- monogalactosyl-diacylglycerol; DGDG- digalactosyl- diacylglycerol; PA- phosphatidic acid; PC- phosphatidylcholine; PE- Phosphatidylethanolamine; PI- Phosphatidylinositol; PS- Phosphatidylserine; PG- Phosphatidylglycerol; TAG- triacyl glycerol; TGDG- trigalactosyl-diacylglycerol.

**Supplementary Figure 3.**
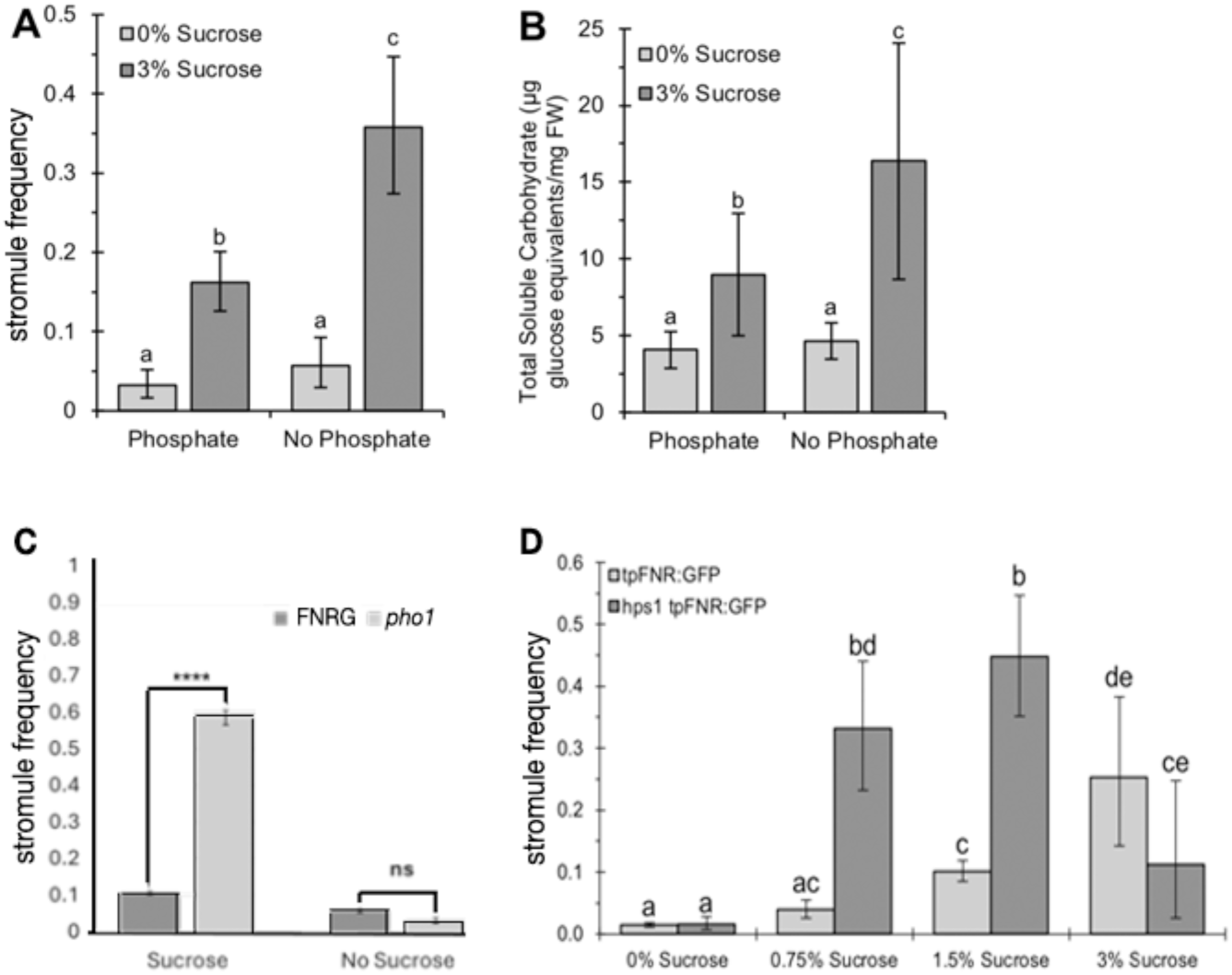
Pi starvation stress (Hartel et al., 2000; 2001; Jouhet et al., 2003) and mutants such as *pho1* (Poirier et al., 1991) and *hps1* (Lei et al., 2011) with increased DGDG exhibit significant increases in stromule frequency on medium supplemented with sucrose. **A.** Phosphate deprivation increases stromule frequency. The hypocotyls of seedlings grown for seven days on media lacking phosphate had similar stromule frequencies to the phosphate containing control for small chloroplasts in epidermal cells, so long as no sucrose was provided in the media (p=0.18, N=20). Phosphate deprivation led to increase in stromule frequency when the media was supplemented with 3% sucrose (p=<0.001, N=20). **B.** Plants grown on media lacking both phosphate and sucrose had total soluble carbohydrate levels comparable to the 0% sucrose control (p=0.70, N=3). The absence of phosphate from the 3% sucrose containing media resulted in significantly higher soluble carbohydrates than the 3% sucrose control (p=0.001, N=3). Error bars represent the 95% confidence interval of the mean. Letters indicate statistically significant groups (p<0.05). N=20 represents two images taken of the upper region of the hypocotyl for each of ten plants. N=3 for carbohydrate analysis represents three tissue samples, each containing the aerial portion of twenty or more seedlings. **C.** The *phosphate* deficient *pho1* mutant (Poirier et al., 1991) has a higher stromule frequency than wild-type. The effect is more pronounced in pho1-tpFNR:GFP relative to tpFNR:GFP when plants are grown on medium with sucrose. Plants of tpFNR-EGFP and *pho1* tpFNR-EGFP were grown on MS medium with 0% or 3% sucrose for 7 days before counting stromule frequency in hypocotyl cells (N=24). Bars represent average stromule frequency ± standard error. Asterisks indicate significant differences within the treatment by a Student’s two tailed t test. **** p < 0.0001. **D.** The *hypersensitive to phosphate starvation1* (*hps1*) mutant overexpresses the SUC2 sucrose importer, that leads to sugar accumulation within the plant (Lei et al., 2011; Lei and Liu, 2011). The mutant transformed with tpFNR-EGFP showed increase in stromule frequencies in small epidermal chloroplasts (sc) in hypocotyls of 7-day-old seedlings with increasing concentrations of sucrose in the media. The *hps1* mutant showed higher frequencies than the control line when sucrose was added, but appeared to reach a point of saturation at the 3% sucrose concentration when large starch grains formed and led to swelling of chloroplasts. Error bars represent the 95% CI of the mean. Letters indicate statistically significant groups (p<0.05). N=8 for stromule frequencies, representing a single image of the upper hypocotyl region in eight plants.

**Supplementary Figure 4.**
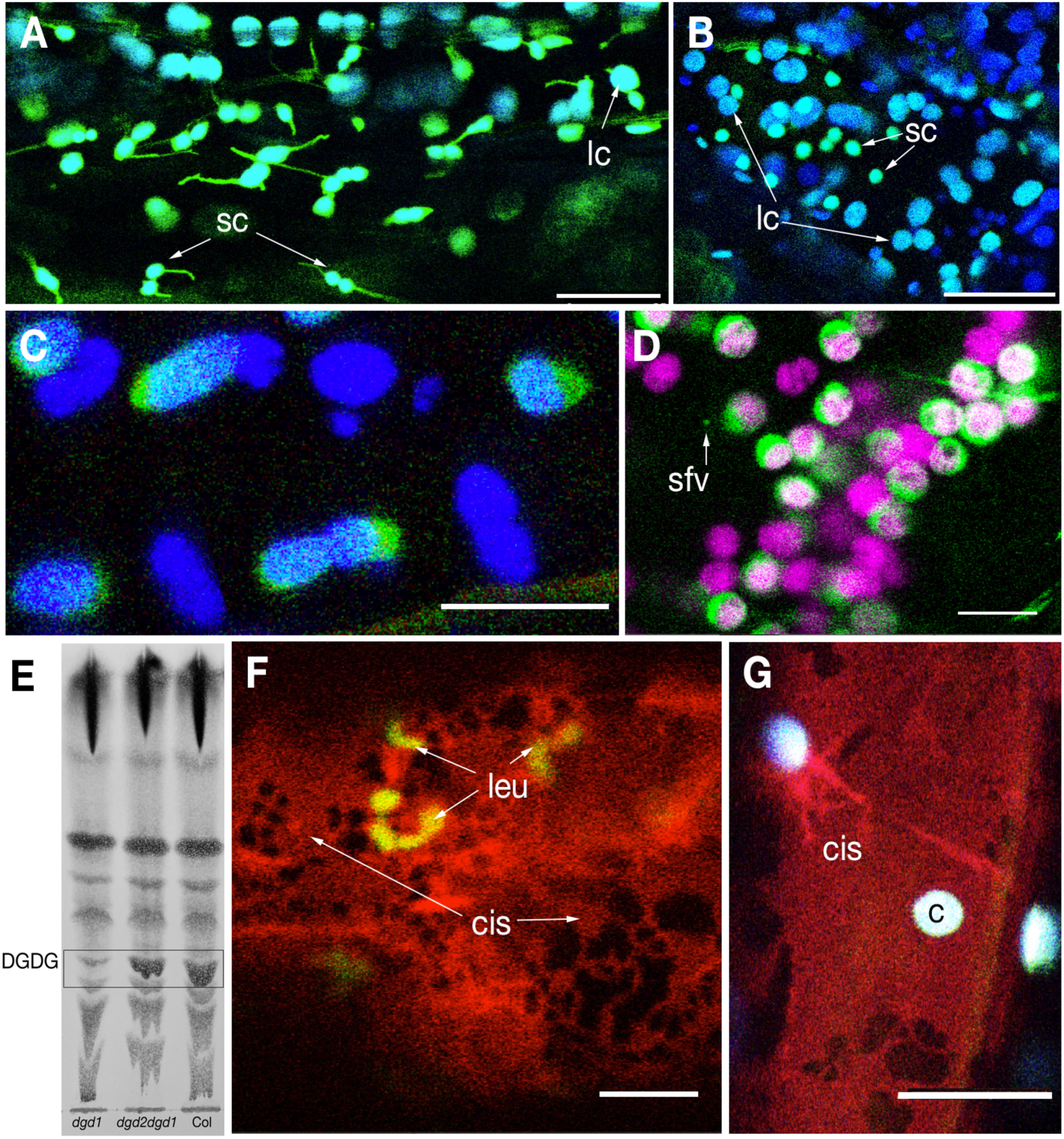
Representative images suggesting that stromule formation in response to 40 mM sucrose treatment is impaired in the *dgd2dgd1* and the *pah1pah2* double mutants with reduced DGDG formation. A. Transgenic line tpFNR-EGFP in Columbia ecotype kept in the dark for 12 hours and then treated with 40 mM sucrose for another 4 hours in the dark shows extended stromules from both small epidermal chloroplasts (sc) and large sub-epidermal chloroplasts (lc). B. In contrast to ‘A’ stromules are not observed from chloroplasts in the *dgd2dgd1* double mutant transgenic for tpFNR-EGFP under the same treatment. C. Representative image showing the swelling up of chloroplasts in the *dgd2dgd1* double mutant transgenic for tpFNR-EGFP after 4 hours of treatment with 40 mM sucrose. Stromules are not observed and the envelope of most chloroplasts appears compromised giving them a fuzzy look. D. A majority of chloroplasts in the the *dgd2dgd1* double mutant transgenic for tpFNR-EGFP after 4 hours of treatment with 40 mM sucrose swell up into spherical forms with a segregation of the stroma (green) and grana (chlorophyll false coloured magenta) compartments. The yinyang chloroplast phenotype indicates a compromised, weak envelope. Stroma filled vesicles (sfv-arrow) released upon plastid breakdown are observed. E. TLC of total lipids extracted from 8 days old *dgd1, dgd2dgd1* double mutant and Columbia wild type seedlings show differences in DGDG levels. F. Epidermal cells exhibit leucoplasts (leu) embedded in the characteristically expanded ER cisternae (cis) in the Arabidopsis *pah1pah2* double mutant expressing stroma-targeted tpFNR-EGFP and RFP targeted to the ER-lumen instead of small chloroplasts observed in wild type (e.g. A). G. Representative image of chloroplasts (c) and expanded ER cisternae (cis) in the *pah1pah2* double mutant expressing tpFNR-EGFP and RFP targeted to the ER suggests lack of stromules following treatment with 40 mM sucrose. Scale bars:; F =5; D,G =10; A, B= 25 µm.

**Supplementary Figure 5.**
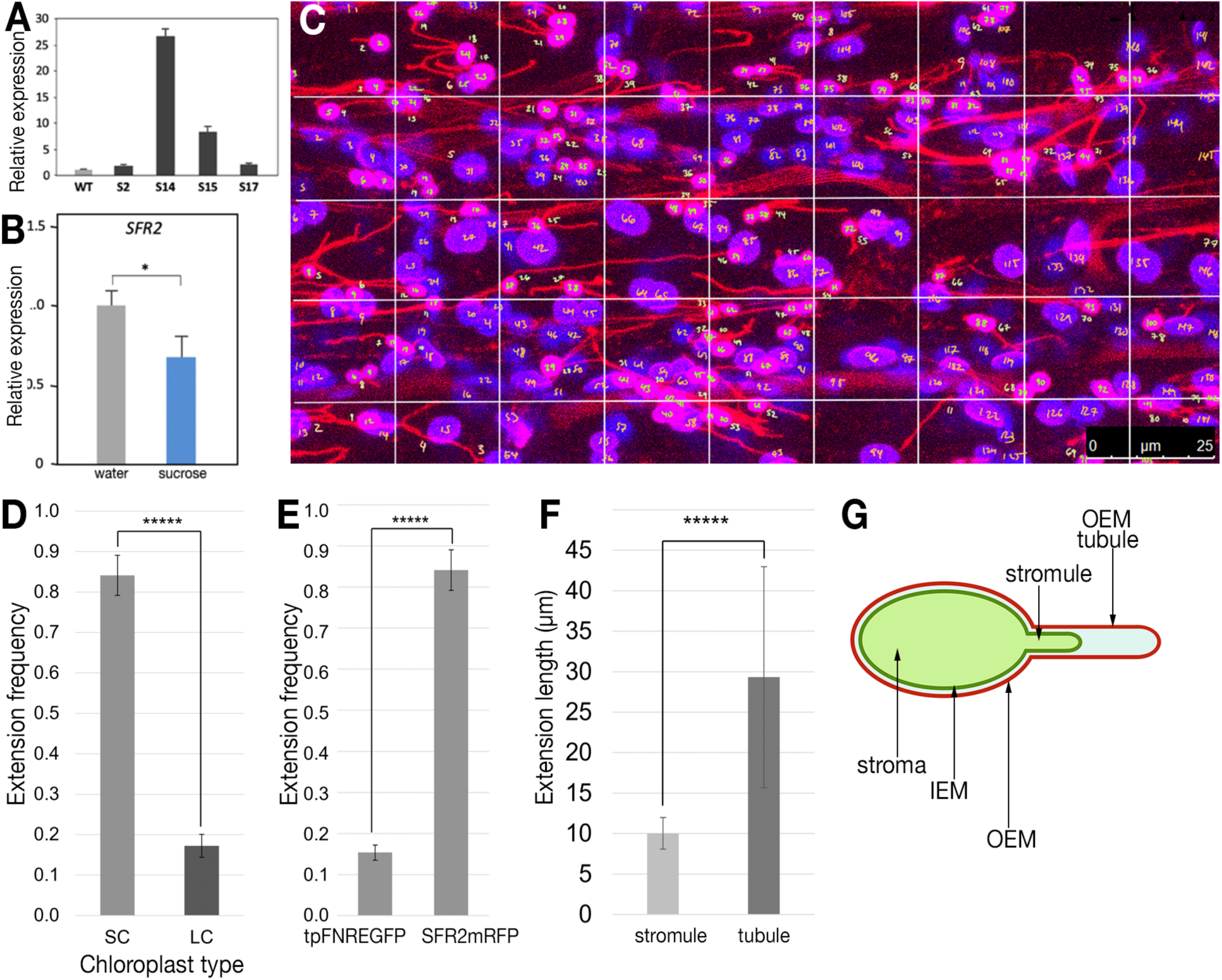
Comparative information on SFR2-mRFP overexpressing lines and a suggested distinction between a stromule and OEM-only tubule. A. PCR based estimation of ***SFR2*** gene expression levels in four independent transgenic lines of pro35S-***SFR2***mRFP shows #14 having the highest overexpression levels. Compared to others the #14 also has a higher number of tubular extensions from chloroplasts. B. Comparing ***SFR2*** expression levels in *Arabidopsis thaliana* (ecotype Columbia) following treatment with water and 40 mM sucrose. The sucrose treatment significantly reduced ***SFR2*** expression. C. D. Small chloroplasts (sc; pink) have a significantly higher stromule frequency than large chloroplasts (lc; appear blue due to chlorophyll fluorescence false colored blue) in sub-epidermal layers. 7-day-old SFR2 mRFP OE soil grown plants were subjected to 18-hour extended period of darkness prior to CLSM imaging for data collection. C – shows a representative image used to count the plastid extensions. D. Bars represent average stromule frequency ± standard error; n = 1500 chloroplasts counted per type. Asterisks indicate significance groups as determined by a students two tailed t-test (p < 0.05). E. Compared to chloroplasts in tpFNR-EGFP plants kept in extended dark period of 18-hours, a significantly large number of plastids in similarly treated SFR2-OE lines exhibited persistent tubular extensions. Bars represent average stromule frequency ± standard error; 8 and 16 biological replicates respectively and n = 1500 chloroplasts counted per type. Asterisks indicate significance groups as determined by a students two tailed t-test (p < 0.05). F. Soil grown, 7-day-old tpFNR-EGFP and SFR2-mRFP (OE line #14) plants observed after 12 hours of darkness showed longer tubular extensions in the SFR2-mRFP line #14. Bars represent average length of extension ± standard deviation; n = 16 chloroplasts with extensions counted per type. Asterisks indicate significance groups as determined by a students two tailed t-test (p < 0.05). G. Diagrammatic depiction of the spatial relation between inner envelope membrane (IEM) and the outer envelope membrane (OEM) suggests distinction between a stromule and an external OEM tubule. The stromule (stroma-filled tubule) lies within an OEM tubule and when bilayer-forming galactolipids increase specifically on the OEM the OEM-tubule may extend beyond the stromule. The idea suggested an explanation for increased tubule persistence at end of dark period (E) and the greater length of tubules (F) extended by plastids in SFR2-mRFP plants.

**Supplementary Figure 6.**
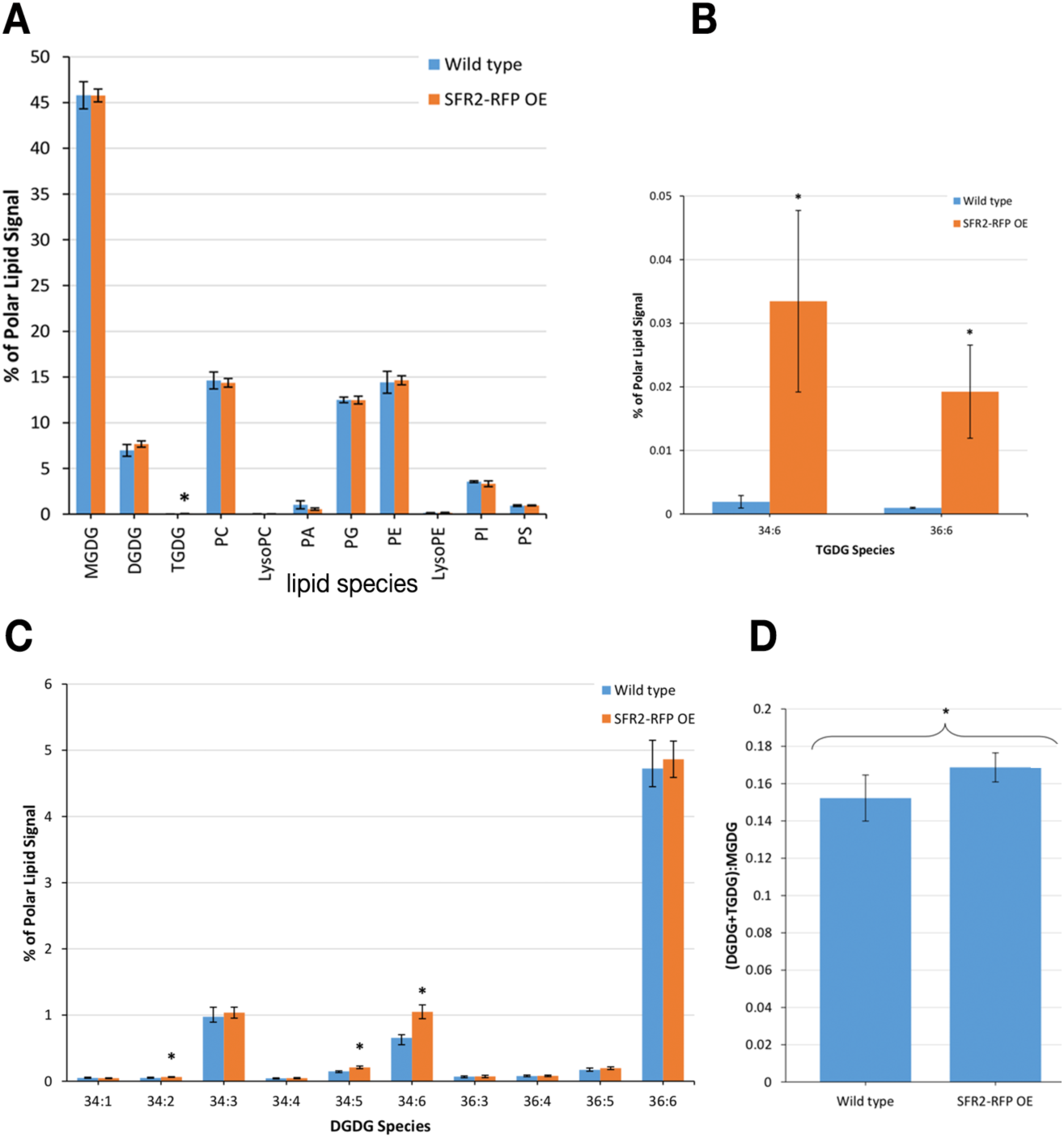
Lipidomics of SFR2-mRFP overexpression line#14. A. Polar lipid content in the *A. thaliana* SFR2-mRFP OE line #14 in comparison to the wildtype. Lipids extracted from rosette leaves of 4 week-old soil-grown plants following overnight darkness were quantified using ESI-MS. Averages ± standard deviation were calculated from five independent samples. Asterisks indicate statistically significant differences between the SFR2 over-expresser and wild type as determined by Student’s t-tests (P < 0.05). The largest difference came in TGDG content of the over-expresser. B. TGDG lipid class content. The average percent polar lipid content of TGDG species 34:6 and 36:6 in the wild type were 0.002 and 0.001 percent, respectively. Under overexpression, the corresponding values increased several folds to 0.033 and 0.019 percent, respectively. C. The average percent polar lipid content of DGDG species 34:1, 34:2, 34:3, 34:4, 34:5, 34:6, 36:3, 36:4, 36:5, and 36:6 in the wild type (blue bars) and under overexpression, (orange bars). According to Student t-tests at the 95% confidence level, 34:2, 34:5, and 34:6 DGDG each had significantly higher content in the over-expresser line (at 0.064, 0.210, and 1.049 percent, respectively) than in the wildtype (at 0.051, 0.150, and 0.656 percent, respectively). D. Average ratio between the lipid content of bilayer-stabilizing galactolipids, DGDG and TGDG, and the non bilayer-forming lipid MGDG in the *A. thaliana* SFR2-RFP OE line #14 with respect to the wildtype. Averages ± standard deviation based on five independent samples suggest statistically significant differences between the over-expresser and the wild type as determined by a Student’s t-test (P < 0.05). Overall, the SFR2-mRFP overexpression line #14 has high bilayer-forming galactolipids. Abbreviations: MGDG, monogalactosyldiacylglycerol; DGDG, digalactosyldiacylglycerol; PA, phosphatidic acid; PC, phosphatidylcholine; PE, Phosphatidylethanolamine; PI, Phosphatidylinositol; PS Phosphatidylserine; PG, Phosphatidylglycerol; TGDG, trigalactosyldiacylglycerol.

**Supplementary Figure 7.**
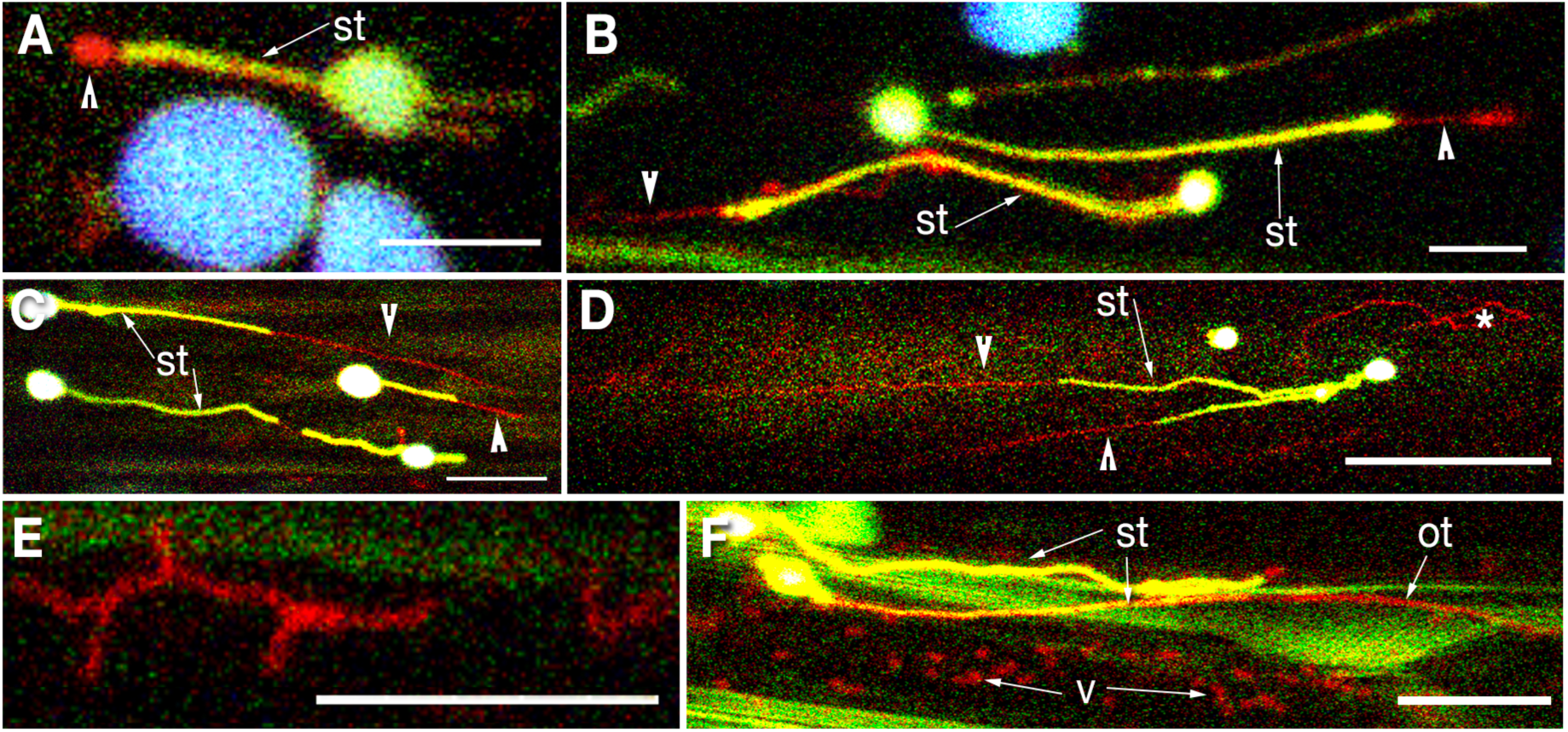
Observations relating to the SFR2-mRFP labelled OEM-tubule that extends beyond the stromule. The plastid OEM lies outside the inner envelope membrane (IEM) that delimits the stroma., stabilized by bilayer-forming galactolipids such as DGDG the OEM may extend beyond the stroma-filled tubule (stromule; st) to form a small protuberance at the tip of a stromule (A; arrowhead) or extend for several micrometres into the cytoplasm to form OEM-only tubules (B, C, D; arrowheads). The OEM-only tubule is dynamic and its length is not fixed Consequently, even in a single snapshot (e.g. C, D) the tips of different stromules may have extensions of different length (arrowheads) or may not any OEM extension at all. Under stress the extra OEM breaks off from the plastid and may form polygonal shapes upon aligning with other cellular membranes (e.g. * in D, E). Eventually the OEM at the tip of a stromule or OEM-blebs break away from the plastids and form independent cytoplasmic vesicles (v; bottom part of F). Size bars: A, B, F = 5 ; C, E =10; D =25 µm.

## Supplementary Movie Legends

**Movie 1. All chloroplasts extend stromules following treatment with 40 mM sucrose in the dark.**

Sixteen consecutive frames from a time-lapse series showing hypocotyl cells of a double transgenic tpFNR-EGFP + RFP-ER plant after 4 hours treatment with 40 mM sucrose in the dark. Notably, both small chloroplasts (sc) in epidermal cells and the relatively large chloroplasts (lc) in sub-epidermal layers exhibit extended stromules. Size bar = 25 µm.

**Movie 2. The OEM extends beyond the IEM-delimited stromule:**

A x,y,t time-lapse series comprising of 42 frames acquired over 5 minutes shows a single epidermal cell chloroplast from a double transgenic N-CHUP-GFP (highlights OEM in green) + CAC1B-mRFP (highlights stroma in red). The outer envelope membrane (green) extended beyond the red stroma. Note that although the tubular plastid extension was clearly subdivided into a stromule and an OEM-only portion, the entire tubule extended and retracted as one. Size bar = 5 µm.

**Movie 3. Stabilized OEM-tubules extending beyond IEM-delimited stromules**

Representative time lapse series taken over 240 seconds showing extra OEM protruding beyond the stroma-filled region in small epidermal chloroplasts in hypocotyl cells soon after 12 hours of darkness. Size bar = 25 µm.

**Movie 4. Asynchronous and independent behaviour of tubules:**

Representative time lapse series taken over 250 seconds from hypocotyl cells of SFR2-mRFP + tpFNR-EGFP double transgenic plants exposed to 110 ± 10 μmol m^-2^ s^-1^ of PAR light after 12 hours of darkness. The extra OEM lying beyond the stroma-filled region in small epidermal chloroplasts starts breaking off to release vesicles into the contiguous cytoplasm. Size bar = 10 µm.

**Movie 5. All chloroplasts extend and shed the OEM:**

Twenty five frames acquired over 75 seconds and played at 5 fps showing OEM-only extension in a small epidermal chloroplast (left, yellow) and a bleb formed at the tip of a, large sub-epidermal chloroplast (right, blue) in hypocotyl cells of a SFR2-mRFP + tpFNR-EGFP double transgenic plant treated with 40 mM sucrose. The series shows that excess DGDG formation results in OEM being shed by all chloroplasts. Red fluorescent SFR2-mRFP-labelled OEM-derived vesicles that have already been shed off into the cytoplasm are also observed. Size bar = 10 µm.

**Movie 6. Stromule extends and retracts within an external OEM tubule:**

Representative time-lapse series taken over 116 seconds from hypocotyl cells of a SFR2-mRFP + tpFNR-EGFP double transgenic shows the dynamic extension and retraction behavior of a green fluorescent stromule within a red-fluorescent OEM tubule. The stromule (yellow, due to red-green overlap) retracts towards the plastid body while remaining within the OEM tubule between frames 12 to 17. A longer intermembrane region is revealed between the retracting stromule and the OEM tubule tip. Re-extension of the stromule is observed within the OEM tubule during frames 24 to 47. The excess OEM remaining beyond the stromule-filled portion of the tubule appears to be shed off during subsequent frames. The time-lapse sequence also underscores the independent behaviour of chloroplasts with respect to OEM-only extensions as a second small chloroplast and the larger sub-epidermal chloroplasts do not exhibit them. Size bar = 5µm.

**Movie 7. Extension of a stromule into an external OEM-tubule:**

A time-lapse series of a small epidermal chloroplast comprising of 14 frames taken over 30 seconds in a hypocotyl cell of a SFR2-mRFP + tpFNR-EGFP double transgenic plant shows the advance of an IEM delimited, stroma-filled region towards the tip of an OEM tubule and fill it up completely. Size bar =10µm.

**Movie 8. Plastid OEM interacts with contiguous ER membranes :**

A x,y,t time-lapse series comprising of 44 frames acquired over 90 seconds from a double transgenic plant expressing OEM -targeted SFR2-mRFP (red) and ER targeted GFP (green) shows the dynamic interactions of the plastid-OEM with the neighbouring ER for two chloroplasts, including the extension of a tubule from the blebbing OEM. Size bar = 10µm.

**Movie 9. Tubular plastid extensions and OEM-derived vesicles enmeshed in the ER:**

A time-lapse x,y,t series of 8 frames acquired over 20 seconds from a hypocotyl cell in a double transgenic line expressing OEM-targeted SFR2mRFP and GFP targeted to the ER shows red fluorescent OEM tubules and vesicles released into the cytoplasm enmeshed in the dynamic ER. Scale bar = 10µm.

**Movie 10. Stromule extending into an OEM-tubule enmeshed in the ER:**

A time-lapse x,y,t series of 35 frames acquired over 88 seconds from a hypocotyl cell in a triple transgenic line expressing stroma-targeted tpFNR-EGFP, OEM targeted SFR2mRFP and GFP targeted to the ER lumen showed the entire tubule (chloroplast on right, stromule and extra OEM-only tubule) enmeshed in and maintaining dynamic behavior in tandem with the neighbouring ER. Over the time course the stromule (yellow) extended into the OEM-only portion (red) to fill it up entirely. Size bar = 10µm.

## Notes

### Competing Interest Statement

The authors have declared no competing interest.

