## Supplementary material for "Tubule extension by plastids is due to increase in bilayer-forming galactolipids at the outer envelope membrane": Single Supplementary file

### Supplementary Table 1: Gene-specific primers used for qPCR

| Primers | Sequence (5′–3′) |
|---|---|
| *Cloning, genotyping* | |
| AtSFR2-Fwd | ATGGAATTATTCGCATTGTTAAT |
| AtSFR2-Rev | GTCAAAGGGTGAGGCTAAAGCA |
| *-qPCR* | |
| AtSFR2-qFwd | GGTGTTCCTGTGCTCGGTTA |
| AtSFR2-qRev | AGTTCGGGCAAGATCATGGG |
| AtDGD1-qFwd | TTGGGAAGCAGCAACACAGA |
| AtDGD1-qRev | ACCACCTCGTTAAAGCTCGG |
| AtDGD2-qFwd | TCGACGCGAATCATCAGCTT |
| AtDGD2-qRev | GCAATGTGTTGCTCCTGCTG |
| AtMGD1-qFwd | AGGTTTCACTGCGATAAAGTGGTT |
| AtMGD1-qRev | AACGGCAATCCCTCCTCAC |
| AtMGD2-qFwd | GATTCGATCACTTCCTATCATCCTC |
| AtMGD2-qRev | TGTGCTAAACCATTCCCCAAC |
| AtMGD3-qFwd | TTGCGGCCTCCCAATTATCC |
| AtMGD3-qRev | GGTGAAAACTCCAGCCCCAT |
| AtACT7-qFwd | ATCCTTGCATCCCTCAGCAC |
| AtACT7-qRev | AACTCACCACCACGAACCAG |
| AtTUB7-qFwd | TTGGGAAGTGGTTAATCT |
| AtTUB7-qRev | GTTGTAGTATACATTAACCCT |

**Supplementary Table 2: Probes used in the present study**

| Name of probe | Characteristic fluorescence | Reference |
|---|---|---|
| tpFNR-EGFP | G- stroma | Schattat et al., 2011 |
| tpFNR-mEosFP | G to R photoconvertible stroma | Schattat et al., 2012 |
| ptyk | Y- stroma | Nelson et al., 2007 |
| ***CAC1***-mRFP | R – stroma | *Choi et al., 1995 |
| N-CHUP1-EGFP | G- OEM | Oikawa et al., 2003 |
| ***OEP7***-GFP | G-OEM, vesicles | Schleiff et al., 2001 |
| ***SFR2***-mRFP | R- OEM | Mathur et al., 2022 |
| ***SFR2***-EGFP | G- OEM | * |
| ***SFR2***-mEosFP | G to R photoconvertible OEM | * |
| tpFNR-EGFP + SFR2-mRFP | G- stroma + R- OEM | * |
| ss- eGFP-HDEL | G- ER lumen | Haseloff et al., 1997 |
| ss-mRFP-HDEL | R – ER lumen | Sinclair et al., 2009 |
| ***SFR2***-mRFP + GFP-ER | R- OEM, G- ER lumen | * |
| ***SFR2***-mRFP + tpFNR-EGFP + GER | R- OEM, G- stroma, G- ER | * |
| ***SFR2***-mRFP + ptyk + GER | R- OEM, Y-stroma, G-ER | * |
| N-CHUP1-EGFP + ***CAC1***-mRFP | G- OEM, R-stroma | * |
| N-CHUP1-GFP + tpFNR-mEosFP | G- OEM, G-R photoconvertible stroma | * |
| *pah1pah2* -tpFNR-EGFP + RER | G- stroma, R-ER lumen | * |

*New probes created.

**Abbreviations:** ***CAC1*** = chloroplastic acetyl-coenzyme A carboxylase AT5G15530 cDNA; mEosFP = monomeric Eos fluorescent protein; *pah1pah2* = phosphatidic acid phosphohydrolase1 x phosphatidic acid phosphohydrolase2 double mutant; ptyk = plastid targeted yellow fluorescent protein; N-***CHUP1*** = N-terminal (aa 1-25) portion of ***CHLOROPLAST UNUSUAL POSITIONING1*** gene; GFP = Green Fluorescent protein; GER = GFP targeted to the ER using a C-terminal lumen retention HDEL peptide; mRFP = monomeric red fluorescent protein; tpFNR-EGFP = transit peptide of ferredoxin-NADP-oxidoreductase (FNR) fused to EGFP ; *tgd1*= *trigalactosyldiacylglycerol1* mutant; SFR2 = SENSITIVE TO FREEZING 2.

**Supplementary Figure 1. Estimation of total soluble carbohydrate levels**

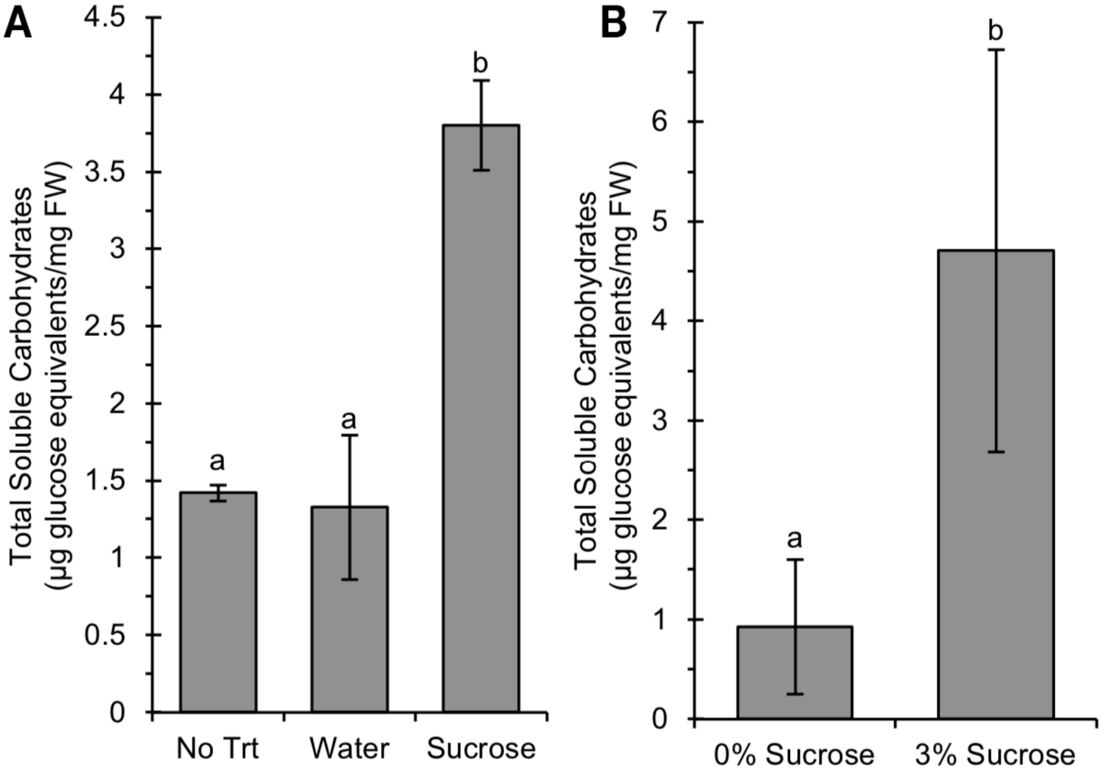

A. 3 h treatment with 40 mM sucrose resulted in higher soluble carbohydrate levels in the above-ground tissue than the water control and no treatment controls (both $p<0.01$, N=4). No difference was seen between the untreated and water treated plants ($p=0.48$, N=4).

B. Plants grown on media containing 3% sucrose had higher total soluble carbohydrate levels than those grown on 0% sucrose media ($p=0.009$, N=3). Error bars indicate the 95% confidence interval of the mean.

For carbohydrate analysis (N= x) represents the number of tissue samples, each containing the aerial portion of thirty seedlings.

**Supplementary Figure 2. Fatty acid and lipid profiles following water and sucrose treatment of tpFNR-EGFP expressing transgenic plants (Columbia ecotype)**

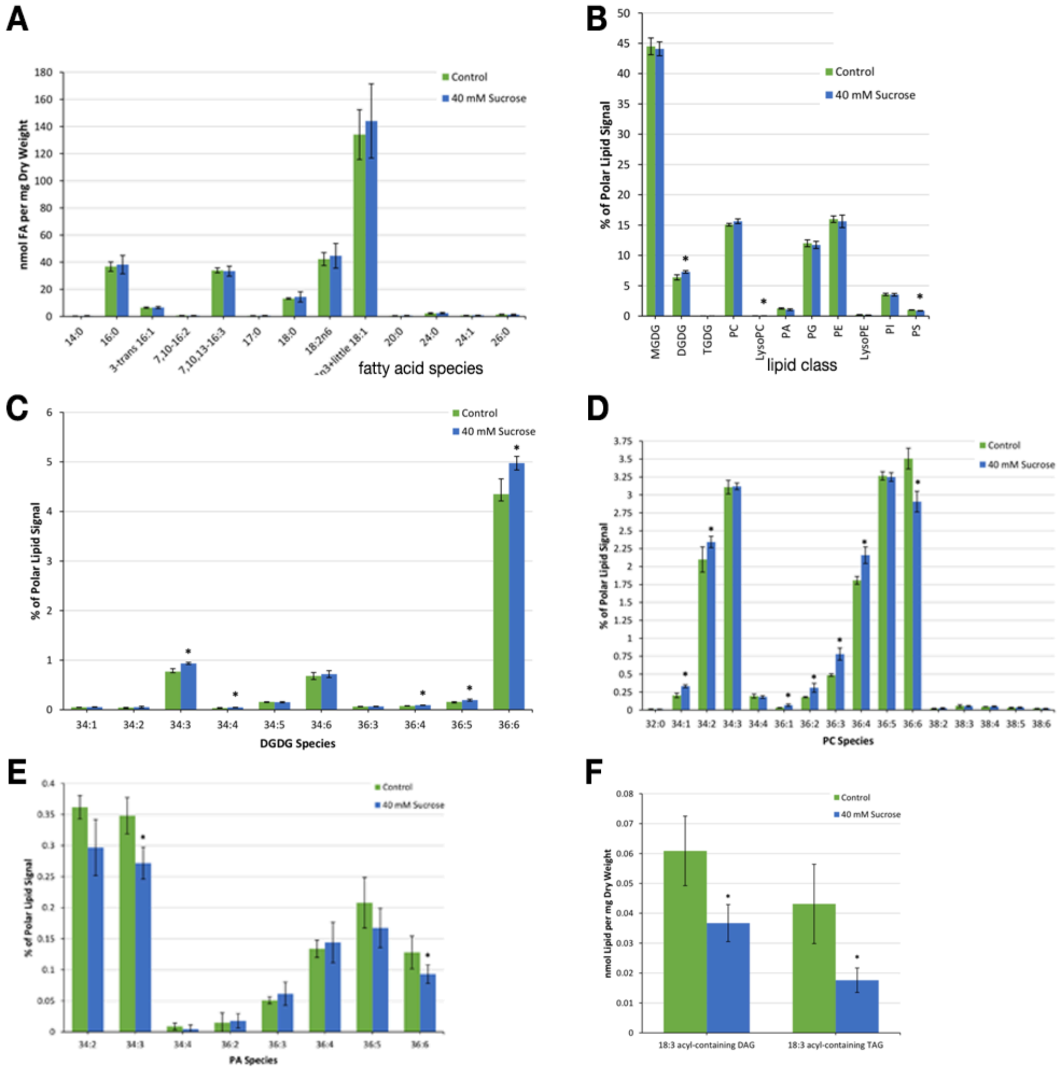

**A.** Comparison of fatty acid profile (in nmol of FA detected per mg dry weight) for *A. thaliana* shows no significant changes following treatment with water and with 40 mM sucrose in the dark for 3 hours. Averages ± standard deviation calculated from five independent samples in 40 mM sucrose treatment and three independent samples in the control. Asterisks indicate statistically significant differences between treatments as determined by Student's t-tests ($P < 0.05$).

**Supplementary Figure 3. Pi starvation stress** (Hartel et al., 2000; 2001; Jouhet et al., 2003) **and mutants such as *pho1*** (Poirier et al., 1991) **and *hps1*** (Lei et al., 2011) **with increased DGDG exhibit significant increases in stromule frequency on medium supplemented with sucrose.**

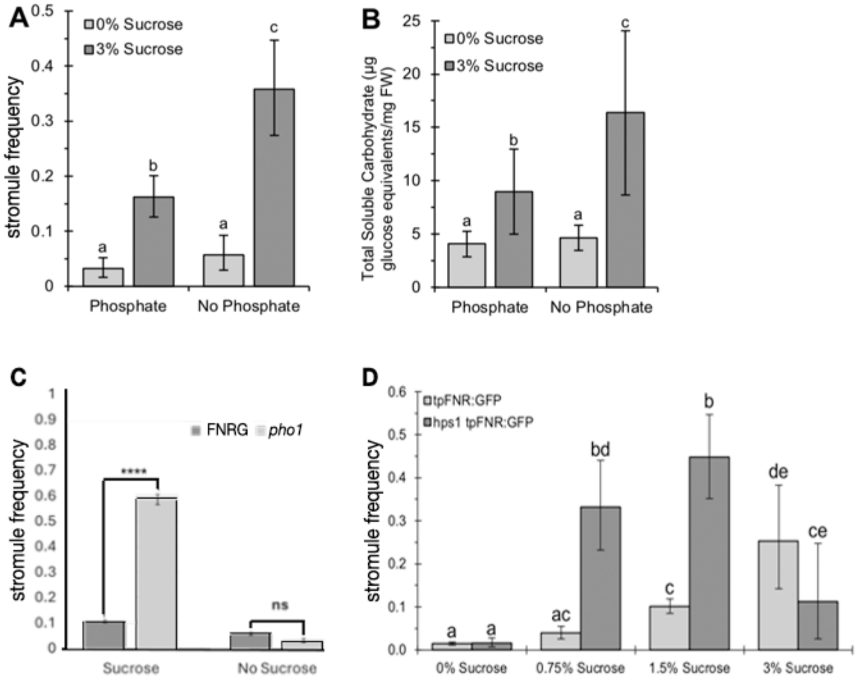

- **A.** Phosphate deprivation increases stromule frequency. The hypocotyls of seedlings grown for seven days on media lacking phosphate had similar stromule frequencies to the phosphate containing control for small chloroplasts in epidermal cells, so long as no sucrose was provided in the media (p=0.18, N=20). Phosphate deprivation led to increase in stromule frequency when the media was supplemented with 3% sucrose (p=<0.001, N=20).
- **B.** Plants grown on media lacking both phosphate and sucrose had total soluble carbohydrate levels comparable to the 0% sucrose control (p=0.70, N=3). The absence of phosphate from the 3% sucrose containing media resulted in significantly higher soluble carbohydrates than the 3% sucrose control (p=0.001, N=3).

Error bars represent the 95% confidence interval of the mean. Letters indicate statistically significant groups (p<0.05). N=20 represents two images taken of the upper region of the hypocotyl for each of ten plants. N=3 for carbohydrate analysis represents three tissue samples, each containing the aerial portion of twenty or more seedlings.

- **C.** The *phosphate* deficient *pho1* mutant (Poirier et al., 1991) has a higher stromule frequency than wild-type. The effect is more pronounced in pho1-tpFNR:GFP relative to tpFNR:GFP when plants are grown on medium with sucrose. Plants of tpFNR-EGFP and *pho1* tpFNR-EGFP were grown on MS medium with 0% or 3% sucrose for 7 days before counting stromule frequency in hypocotyl cells (N=24). Bars represent average stromule frequency ± standard error. Asterisks indicate significant differences within the treatment by a Student's two tailed t test. **** p < 0.0001.

**D.** The *hypersensitive to phosphate starvation1* (*hps1*) mutant overexpresses the SUC2 sucrose importer, that leads to sugar accumulation within the plant (Lei et al., 2011; Lei and Liu, 2011). The mutant transformed with tpFNR-EGFP showed increase in stromule frequencies in small epidermal chloroplasts (sc) in hypocotyls of 7-day-old seedlings with increasing concentrations of sucrose in the media. The *hps1* mutant showed higher frequencies than the control line when sucrose was added, but appeared to reach a point of saturation at the 3% sucrose concentration when large starch grains formed and led to swelling of chloroplasts. Error bars represent the 95% CI of the mean. Letters indicate statistically significant groups ($p<0.05$). N=8 for stromule frequencies, representing a single image of the upper hypocotyl region in eight plants.

**Supplementary Figure 4. Representative images suggesting that stromule formation in response to 40 mM sucrose treatment is impaired in the *dgd2dgd1* and the *pah1pah2* double mutants with reduced DGDG formation.**

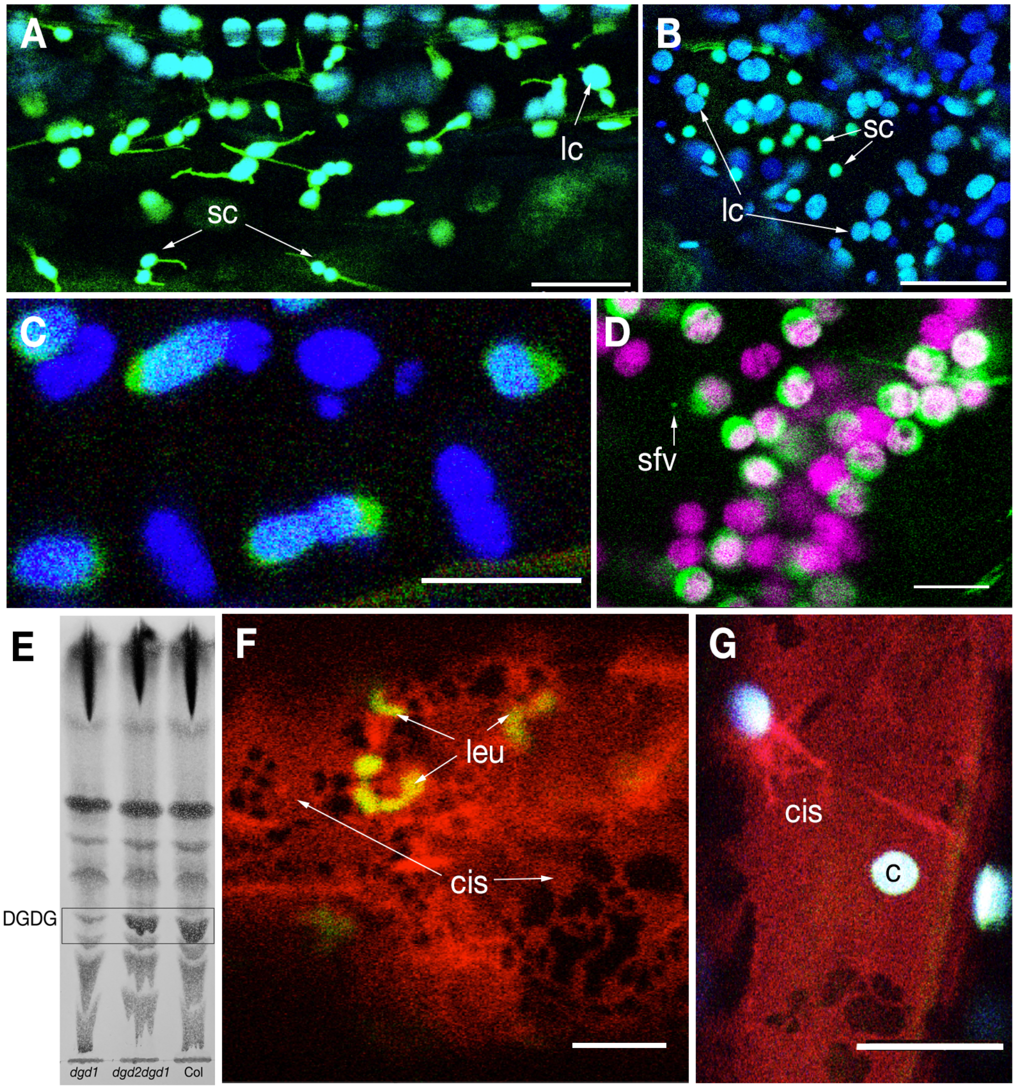

A. Transgenic line tpFNR-EGFP in Columbia ecotype kept in the dark for 12 hours and then treated with 40 mM sucrose for another 4 hours in the dark shows extended stromules from both small epidermal chloroplasts (sc) and large sub-epidermal chloroplasts (lc).

B. In contrast to 'A' stromules are not observed from chloroplasts in the *dgd2dgd1* double mutant transgenic for tpFNR-EGFP under the same treatment.

C. Representative image showing the swelling up of chloroplasts in the *dgd2dgd1* double mutant transgenic for tpFNR-EGFP after 4 hours of treatment with 40 mM sucrose. Stromules are not observed and the envelope of most chloroplasts appears compromised giving them a fuzzy look.

D. A majority of chloroplasts in the the *dgd2dgd1* double mutant transgenic for tpFNR-EGFP after 4 hours of treatment with 40 mM sucrose swell up into spherical forms with a segregation of the stroma (green) and grana (chlorophyll false coloured magenta) compartments. The yinyang chloroplast phenotype indicates a compromised, weak envelope. Stroma filled vesicles (sfv-arrow) released upon plastid breakdown are observed.

E. TLC of total lipids extracted from 8 days old *dgd1, dgd2dgd1* double mutant and Columbia wild type seedlings show differences in DGDG levels.

F. Epidermal cells exhibit leucoplasts (leu) embedded in the characteristically expanded ER cisternae (cis) in the Arabidopsis *pah1pah2* double mutant expressing stroma-targeted tpFNR-EGFP and RFP targeted to the ER-lumen instead of small chloroplasts observed in wild type (e.g. A).

G. Representative image of chloroplasts (c) and expanded ER cisternae (cis) in the *pah1pah2* double mutant expressing tpFNR-EGFP and RFP targeted to the ER suggests lack of stromules following treatment with 40 mM sucrose.

Scale bars:; F =5; D,G =10; A, B= 25 µm.

**Supplementary Figure 5. Comparative information on SFR2-mRFP overexpressing lines and a suggested distinction between a stromule and OEM-only tubule.**

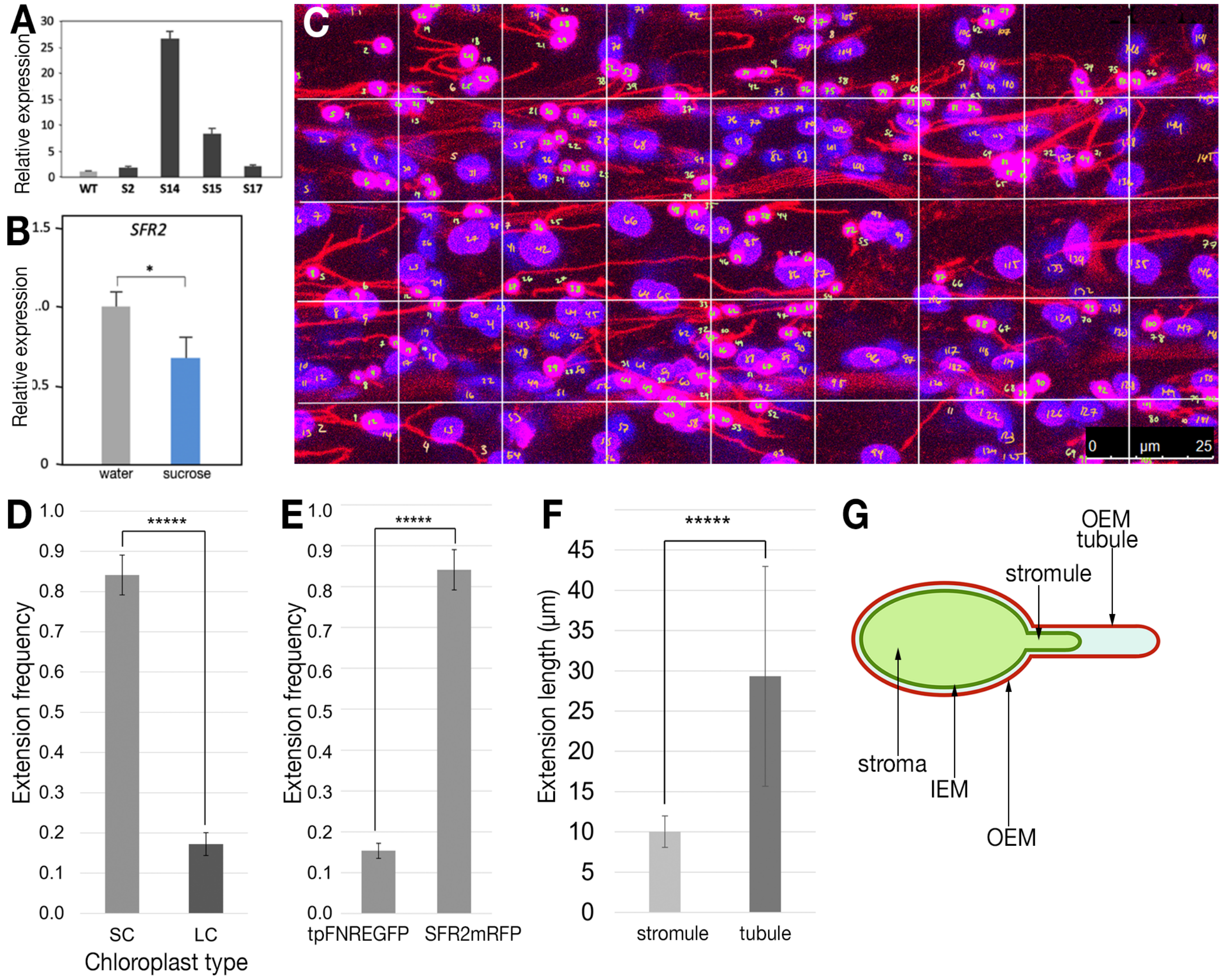

A. PCR based estimation of ***SFR2*** gene expression levels in four independent transgenic lines of pro35S-***SFR2***mRFP shows #14 having the highest overexpression levels. Compared to others the #14 also has a higher number of tubular extensions from chloroplasts.

**Supplementary Figure 6. Lipidomics of SFR2-mRFP overexpression line#14**

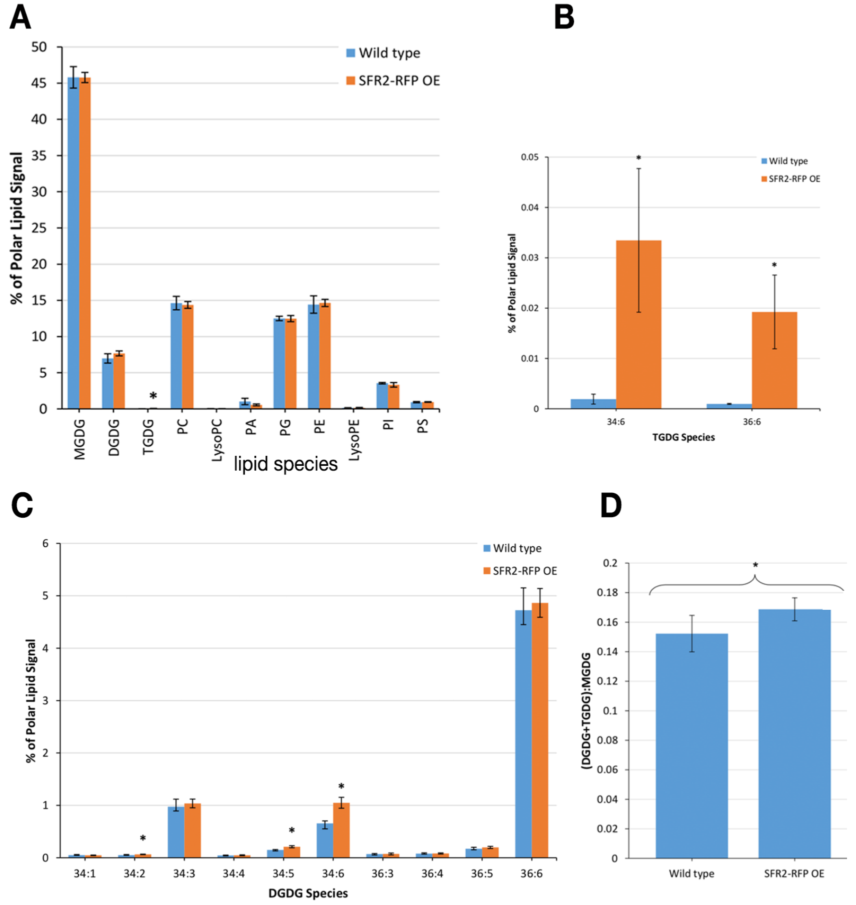

A. Polar lipid content in the *A. thaliana* SFR2-mRFP OE line #14 in comparison to the wildtype. Lipids extracted from rosette leaves of 4 week-old soil-grown plants following overnight darkness were quantified using ESI-MS. Averages ± standard deviation were calculated from five independent samples. Asterisks indicate statistically significant differences between the SFR2 over-expresser and wild type as determined by Student's t- tests ($P < 0.05$). The largest difference came in TGDG content of the over-expresser.

B. TGDG lipid class content. The average percent polar lipid content of TGDG species 34:6 and 36:6 in the wild type were 0.002 and 0.001 percent, respectively. Under overexpression, the corresponding values increased several folds to 0.033 and 0.019 percent, respectively.

C. The average percent polar lipid content of DGDG species 34:1, 34:2, 34:3, 34:4, 34:5, 34:6, 36:3, 36:4, 36:5, and 36:6 in the wild type (blue bars) and under overexpression, (orange bars). According to Student t-tests at the 95% confidence level, 34:2, 34:5, and 34:6 DGDG each had significantly higher content in the over-expresser line (at 0.064, 0.210, and 1.049 percent, respectively) than in the wildtype (at 0.051, 0.150, and 0.656 percent, respectively).

D. Average ratio between the lipid content of bilayer-stabilizing galactolipids, DGDG and TGDG, and the non bilayer-forming lipid MGDG in the *A. thaliana* SFR2-RFP OE line #14 with respect to the wildtype. Averages ± standard deviation based on five independent samples suggest statistically significant differences between the over-expresser and the wild type as determined by a Student's t-test ($P < 0.05$). Overall, the SFR2-mRFP overexpression line #14 has high bilayer-forming galactolipids.

Abbreviations: MGDG, monogalactosyldiacylglycerol; DGDG, digalactosyldiacylglycerol; PA, phosphatidic acid; PC, phosphatidylcholine; PE, Phosphatidylethanolamine; PI, Phosphatidylinositol; PS Phosphatidylserine; PG, Phosphatidylglycerol; TGDG, trigalactosyldiacylglycerol.

**Supplementary Figure 7. Observations relating to the SFR2-mRFP labelled OEM-tubule that extends beyond the stromule.**

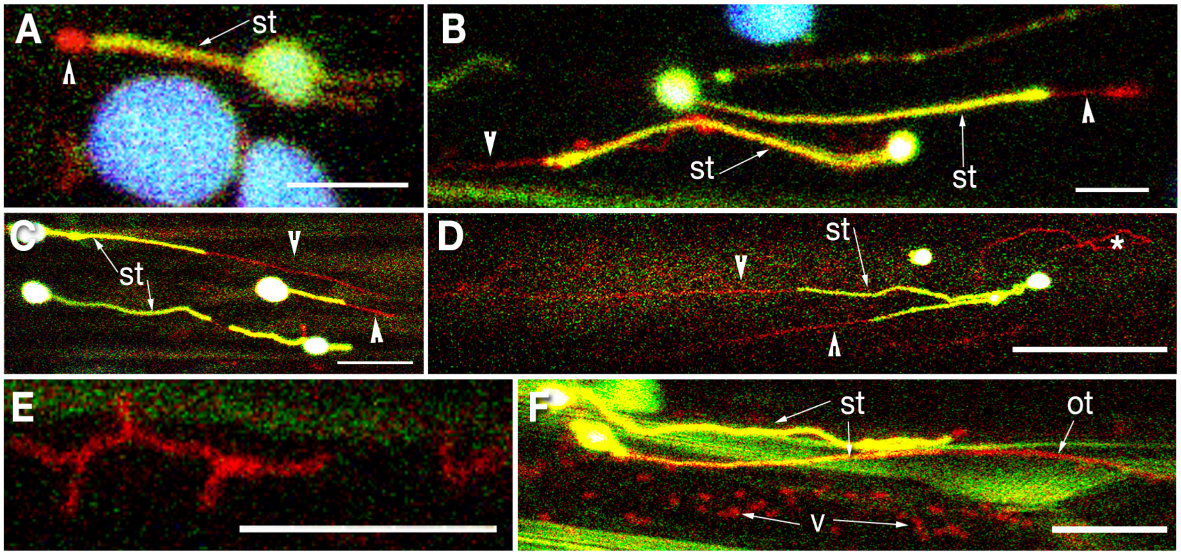

The plastid OEM lies outside the inner envelope membrane (IEM) that delimits the stroma., stabilized by bilayer-forming galactolipids such as DGDG the OEM may extend beyond the stroma-filled tubule (stromule; st) to form a small protuberance at the tip of a stromule (A; arrowhead) or extend for several micrometres into the cytoplasm to form OEM-only tubules (B, C, D; arrowheads). The OEM-only tubule is dynamic and its length is not fixed Consequently, even in a single snapshot (e.g. C, D) the tips of different stromules may have extensions of different length (arrowheads) or may not any OEM extension at all. Under stress the extra OEM breaks off from the plastid and may form polygonal shapes upon aligning with other cellular membranes (e.g. * in D, E). Eventually the OEM at the tip of a stromule or OEM-blebs break away from the plastids and form independent cytoplasmic vesicles (v; bottom part of F). Size bars: A, B, F = 5 ; C, E =10; D =25 μm.
